# Deciphering the Dynamics of Leukemogenic Transformation from Clonal Hematopoiesis in Congenital Neutropenia

**DOI:** 10.64898/2026.11.10.765578

**Authors:** M. Klimiankou, S. Kandabarau, C. Zeidler, S. Mellor-Heineke, M. Haar, Y. L. Behrens, D. C. Dale, V. Makaryan, M. Kelley, A. A. Bolyard, C. M. Niemeyer, M. Erlacher, A. Yoshimi-Nöllke, P. E. Newburger, M. Mezger, P. Lang, T. Haferlach, N. Nadarajah, C. Lengerke, K. Welte, J. Skokowa

## Abstract

Inherited neutropenia syndromes, severe congenital neutropenia (CN) and Shwachman-Diamond syndrome (SDS), confer a high cumulative risk of MDS and AML. It is not yet clear which somatic mutations, in which combination, allele frequency and sequence identify patients at high risk. We analyzed clonal hematopoiesis (CH) in the largest representative cohort of inherited neutropenia patients to date - 658 samples from 309 patients - using error-corrected panel sequencing with longitudinal sampling. This was complemented in selected cases by single-cell DNA sequencing and genotyping of single-cell derived colonies. CH was detected in early childhood, affecting 37.7% of CN and 60% of SDS patients. The somatic mutation burden remained essentially unchanged over decades. In CN, *CSF3R* mutations were dominant across genetic subtypes, while additional mutations were rare and included *ASXL1*, *TET2*, and *DNMT3A*. *ASXL1* mutations most frequently co-occurred with *CSF3R* mutations. Co-occurring *CSF3R* and *TP53* mutations were only found in *SRP54*-CN and *CLPB*-CN patients. In SDS, *TP53* was the most frequently mutated gene, followed by low-prevalence somatic mutations in *DNMT3A*, *IDH1*, *PTPN11*, and *RUNX1*. In CN-MDS/AML*, RUNX1* and *SETBP1* mutations emerged shortly before progression to MDS/AML, and membrane-proximal *CSF3R* mutations only at the leukemic stage. In SDS, biallelic *TP53*-mutant clones expanded prior to transformation, and a variant allele frequency (VAF) of dominant *TP53* clone above 0.074 discriminated between patients who progressed and those who did not. Peripheral blood reflected bone marrow genotype for most key drivers. Leukemic transformation in inherited neutropenia is therefore not stochastic, but rather follows a defined mutational sequence, initiated in childhood.

## Introduction

Patients with pre-leukemia bone marrow failure syndromes (BMFS) frequently progress to myelodysplastic syndrome (MDS) or leukemia, mainly acute myeloid leukemia (AML)^1,2^. The inherited mutations underlying BMFS impose intrinsic and extrinsic stress on hematopoietic stem and progenitor cells (HSPCs), predisposing them to leukemogenic transformation at a young age^3–5^. Transformation is preceded by the emergence, persistence, and expansion of HSPC clones carrying mutations in leukemia-associated genes, a process known as clonal hematopoiesis (CH)^6–8^. In BMFS patients, CH is often detectable in childhood^9^, in contrast to healthy individuals, in whom CH typically appears late in life^1,6,8^. Distinct BMFSs are characterized by specific patterns of CH-associated mutations, with some shared genes. *RUNX1, ASXL1, STAG2, SETBP1* mutations have been reported in patients with GATA2 deficiency, SAMD9/SAMD9L syndromes, and severe congenital neutropenia (CN)^1,10–15^, while *TP53* mutations are the most common in Shwachman–Diamond syndrome (SDS)^3–5,14–16^. Age-related CH is dominated by *DNMT3A* and *TET2* mutations^6–8^. Cytogenetic abnormalities are often early and dominant events preceding leukemogenic transformation in certain BMFS, such as monosomy 7 and trisomy 8 in GATA2 or SAMD9/SAMD9L syndromes, and the gain of 1q and 3q, in addition to monosomy 7, in Fanconi anemia^17^.

Patients with CN or cyclic neutropenia (CyN) experience early-onset, life-threatening infections and are treated with lifelong daily injections of granulocyte colony-stimulating factor (G-CSF)^18^. They frequently acquire truncating mutations in *CSF3R*, which encodes the G-CSF receptor (G-CSFR)^9,19–21^ and their cumulative risk of MDS or AML is approximately 20%^19,22,23^. The number and variant allele frequencies (VAF) of *CSF3R-*mutant hematopoietic clones vary between patients and may fluctuate over time within the same patient. Unlike *CSF3R* mutations, acquisition of *RUNX1* mutations is often associated with rapid progression to MDS or AML^10,12^. However, not all CN or CyN patients who develop MDS or AML acquired *RUNX1* or even *CSF3R* mutations. SDS patients may have severe or mild neutropenia, thrombocytopenia, or anemia^16^. In these patients, CH involves acquiring *TP53* mutations, and while *TP53*-mutant clones can persist for years without transformation, biallelic *TP53* mutations can trigger rapid leukemogenic progression^16,24,25^.

A deep understanding of the mutational landscape preceding leukemogenic transformation is essential for guiding clinical decisions, including timely hematopoietic stem cell transplantation (HSCT). Predicting which patients will transform requires knowledge not only about which acquired mutations patients have, but in which order, in which clones, and how long before overt disease they can be detected. To develop such robust predictive algorithms of a rare patient population, it is crucial to analyze a broad set of leukemia-associated genes with high sensitivity and specificity in large patient cohorts. Single-cell sequencing can further deepen our understanding of leukemogenic evolution in hematopoietic cells caused by specific mutations and their combinations. Such insights will also help to define the mechanisms underlying the emergence of CH and its evolution to leukemia, supporting the development of therapeutic strategies to prevent CH and leukemogenic progression. Furthermore, understanding the sensitivity of CH detection in bone marrow (BM) compared with peripheral blood (PB) is important for clinical diagnostics.

Here we analyze CH and its evolution to MDS/AML in the largest cohort of CN, CyN, and SDS patients reported to date, combining duplex-UMI error-corrected panel sequencing with longitudinal sampling, single-cell DNA sequencing and genotyping of single-cell-derived colonies. We define the sequence in which driver mutations are acquired, the interval over which they are detectable before transformation, and the sensitivity of mutation detection in BM compared with PB.

## Material and Methods

### Human subjects

PB and BM samples were acquired from CN, SDS patients, and healthy donors (HD) under informed consent approved by the Ethics Committee of the University Hospital Tübingen. The same applies to cord blood and PB samples donated by healthy donors.

### Study cohort characteristics

A total of 756 samples derived from four distinct cohorts were analyzed. The primary cohorts consisted of 281 patients with CN and SDS who did not develop leukemia and 37 CN and SDS patients who subsequently progressed to MDS/AML (CN-MDS/AML, SDS-MDS/AML, respectively). The study included 20 peripheral blood (PB) samples from healthy donors of varying ages to facilitate a comparison between age-associated and CN-driven CH. A Panel of Normals (PoN), consisting of cord blood samples from 20 donors, was utilized to filter out panel-specific and sequencing artifacts (**Table S1**). Out of 318 patients included in the study, 283 had a genetic diagnosis of CN or SDS (**Table S2**) and 35 were genetically unclassified after whole exome or genome sequencing but met inclusion criteria based on persistent neutropenia starting at birth, and/or family history of neutropenia. Patients with autoimmune neutropenia were not included in the study. The median patient age at the time of the first sample collection was 10.7 years (range 17 days -65.12 years), 166 were males (52%) and 152 females (48%). For 169 patients, sequential samples were sequenced with a median observation interval of 2.91 years (observation range of 27 days - 24.6 years).

The genetic groups covered the entire spectrum of neutropenia-associated genetic entities, including 122 *ELANE (41 ELANE*-CN*, 32 ELANE-*CyN and 49 *ELANE*-CN/CyN), 61 *SBDS*, 38 *HAX1*, 14 *SRP54*, 10 *G6PC3*, 7 *JAGN1*, 5 *CLPB* patients, rare genetic groups consisting of 3 patients (*TAFAZZIN*, *TCIRG1*, *GFI1*, *WAS*, *SLC37A4*, *VPS13B),* and 1-2 patients (*CXCR4*, *ADA2*, *AP1B1*, *CSF3R*, *CXCR2*, *GINS1*, *SEC61A1*) (**Figure 1A, Table S2**). Of the 37 CN-MDS/AML and SDS-MDS/AML patients, 9 (5 *ELANE*, 2 *HAX1*, 1 *SBDS*, and 1 *SLC37A4*) lacked samples prior to leukemia and were therefore excluded from the CH analysis. Furthermore, 10 patients did not have samples from the MDS/AML stage, restricting their available data to either samples prior to leukemia (n=7) or post-BMT/relapse samples (n=3) (**Table S1**).

**Figure 1.**
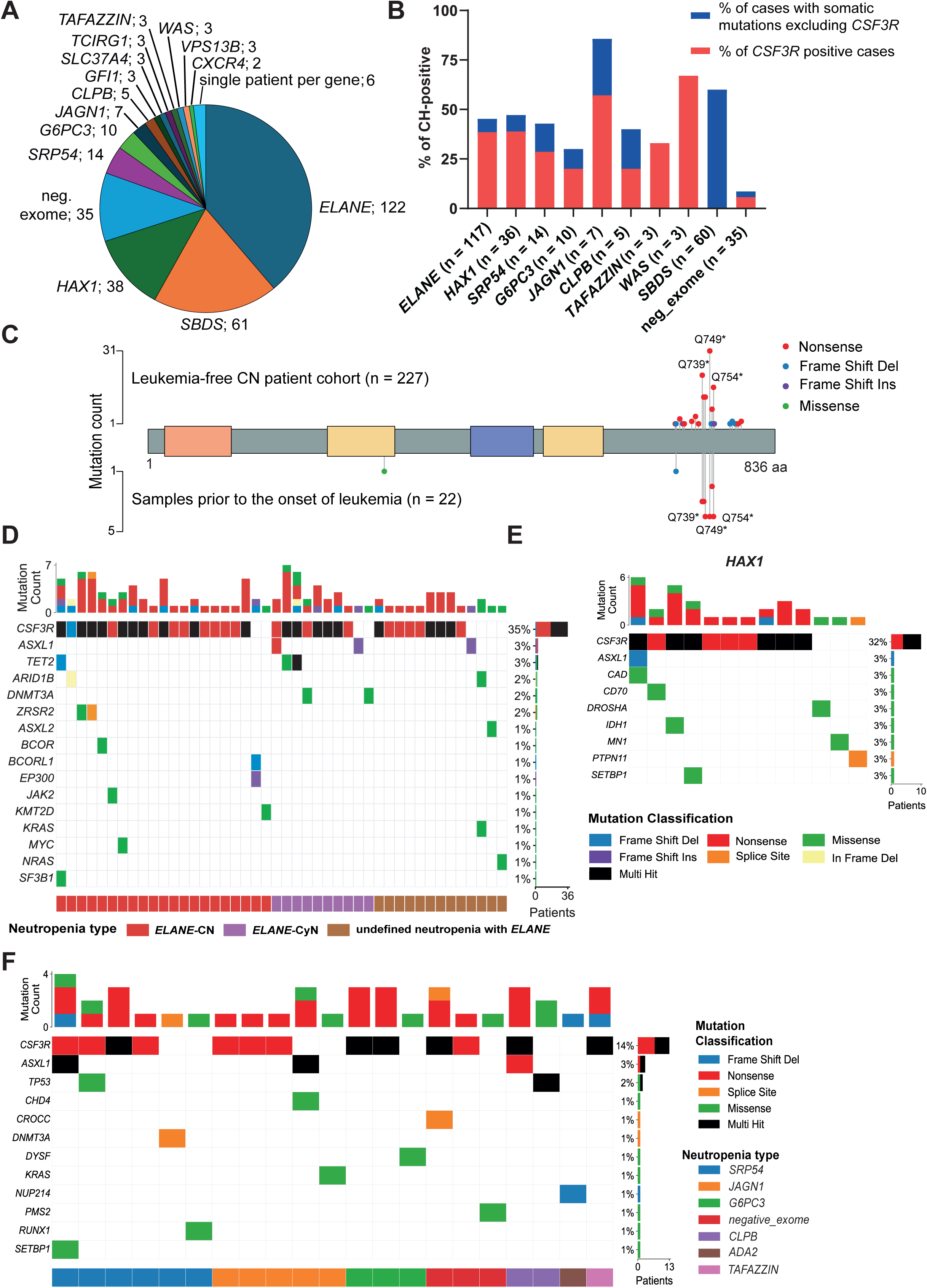
Mutational landscape and prevalence of CH across genetic subgroups of congenital neutropenia patients. **A,** The distribution of patients across specific germline genetic entities. The “single patient per gene” group consists of subjects with *ADA2, AP1B1, CSF3R, CXCR2, GINS1, SEC61A1* mutations. **B,** Prevalence of CH across CN genetic subgroups, *SBDS* patients, and genetically unclassified (neg_exome) cases examined at neutropenia stage. Stacked bar charts represent the percentage of patients in each genetic category who tested positive for CH. Red indicates cases harboring somatic *CSF3R* mutations, and blue represents cases with somatic mutations in any of the other leukemia-associated genes analyzed, but no mutations in *CSF3R*. **C,** Lollipop plot illustrates the frequencies of *CSF3R* mutations in leukemia-free CN patients compared to samples from CN-MDS/AML patients prior leukemia. Mutation types are color-coded (nonsense, frame shift del, frame shift ins, missense). The three most frequent *CSF3R* mutations are annotated. **D-F**, The spectrum of somatic mutations in leukemia-free 33 *ELANE*-CN, 28 *ELANE*-CyN, 43 undefined neutropenia (CN or CyN) with *ELANE* mutations (**D**), 31 *HAX1* (**E**), and rare (**F**) genetic groups. Top bar plots indicate individual mutation counts per patient. Each column represents a CN patient. Neutropenia types and mutation classifications are color-coded according to the legend key. Only patients with somatic mutations are shown. Abbreviations: CN, congenital neutropenia; CyN, cyclic neutropenia; VAF, variant allele frequency.

### Design of capture panels for the detection of somatic leukemia-associated mutations

Using publicly available data (https://www.cbioportal.org/) on sequencing of myeloid neoplasm (n = 2613), we extracted the genomic coordinates of acquired mutations with positive samples higher than 2. The exon coordinates of genes that are recurrently mutated in CN-MDS/AML were added to the panel. The resulting bed file was used to design and synthesize 2571 hybridization capture probes by Integrated DNA Technologies. Integration of UMIs in sequencing reads enables strand-specific molecular indexing and thus sequencing error correction. The list of genes and gene regions covered by capture panel version 1 is shown in **Table S3**.

The same technology was also utilized to produce capture probes for panel version 2. This revised panel was size optimized based on data generated using panel version 1. Capture panel version 2 consists of 1275 capture probes that cover entire coding regions of 29 genes recurrently mutated in leukemia and hot-spot regions in 16 genes (**Table S4**).

## Results

### Prevalence of clonal hematopoiesis across patient genotypes

The study included 318 patients whose samples were available through the research laboratory of the Severe Chronic Neutropenia International Registry (SCNIR). We assessed CH in 309 patients, because for the remaining nine patients only leukemic samples were available for the study. CH was detected in 42.1% (130/309) of patients from the combined CN and SDS cohorts analyzed at the neutropenia stage in the absence of leukemogenic transformation. CH prevalence was highest in *JAGN1* patients (85.7%, 6/7), followed by *SDS* (60%, 36/60), *HAX1* (47.2%, 17/36), *ELANE* (45.3%, 53/117), *SRP54* (42.9%, 6/14), and *CLPB* group (40%, 2/5) (**Figure 1B**). Among *ELANE-*mutated patients, CH was more frequent in *ELANE*-CN (51.2%, 21/41) than in *ELANE*-CyN (31.3%, 10/32). Two of the three *WAS* patients were CH-positive. CH rates were lower in the *G6PC3* (30%, 3/10) and *TAFAZZIN* (33.3%, 1/3) groups (**Figure 1B**), and CH was detected in the single *ADA2* patient. CH was considerably less frequent in genetically unclassified neutropenia patients (8.6%, 3/35). Applying the clonal hematopoiesis of indeterminate potential (CHIP)-defining threshold of VAF > 2%^26^ reduced CH prevalence most in *HAX1* and SDS, whereas CH in *WAS*, *TAFAZZIN* and *CLPB* consisted entirely of clones above this threshold (**Figure S1A**).

Of 75 CN patients with *CSF3R* mutations, 34 (45%) carried two or more (range 2 -6) mutations (**Figure S1B**). The most frequent variants were p.Q749* (31 patients), followed by p.Q739* (21 patients) and p.Q754* (16 patients). In samples from patients who later developed leukemia (prior to leukemia), *CSF3R* mutations clustered between residues Q739 and Q754, whereas in patients who remained leukemia-free they were more widely distributed, ranging from p.E703* to p.G791* (**Figure 1C**). *CSF3R* mutations were more frequent in *ELANE*-CN (19 out of 33) than in *ELANE*-CyN (8 out of 28) (OR 3.32, 95% CI 1.03-11.5, *p*-value 0.038) (**Figure 1D**). Other somatic mutations in leukemia-free CN patients with *ELANE*, *HAX1*, *SRP54, JAGN1*, *G6PC3,* and *CLPB* were rare (**Figure 1D-F**). *CSF3R* and *TP53* mutations co-occurred in individual *CLPB-*and *SRP54*-CN patients (**Figure 1F, S1C**).

*ASXL1* mutations, frameshift and nonsense, were the most frequent after *CSF3R*. Thus, acquired *ASXL1* mutations were detected in 10 of 249 CN patients at the neutropenic stage (4.0%). Of the patients with *ASXL1* mutations, seven also carried *CSF3R* mutations (**Figure 2A-E, S1D-E**). In at least two leukemia-free CN patients, *CSF3R* and *ASXL1* mutations were present in the same clone (**Figure 2C-D**). In an *SRP54* patient, we additionally detected a *SETBP1* p.G870S mutation at low VAF; this mutation was not detected by scDNA-seq. Mutations in age-related CH genes were rare in CN: *DNMT3A* in 7 (2.8%) and *TET2* in 6 (2.4%) of 249 patients. One 45-year-old *ELANE*-CyN patient had *TET2*, *DNMT3A*, *CSF3R* and three *ASXL1* mutations (**Figure 2E**). Three of ten CN patients with *ASXL1* mutations at the neutropenia stage developed MDS/AML. In two of them, leukemia-initiating clones already carried both *CSF3R* and *ASXL1* mutations more than two years before overt MDS/AML (**Figure S1D-E**). Conversely, *ASXL1*-mutant CH persisted without progression for up to 6.75 years of follow-up in another patient (**Figure S1F**).

**Figure 2.**
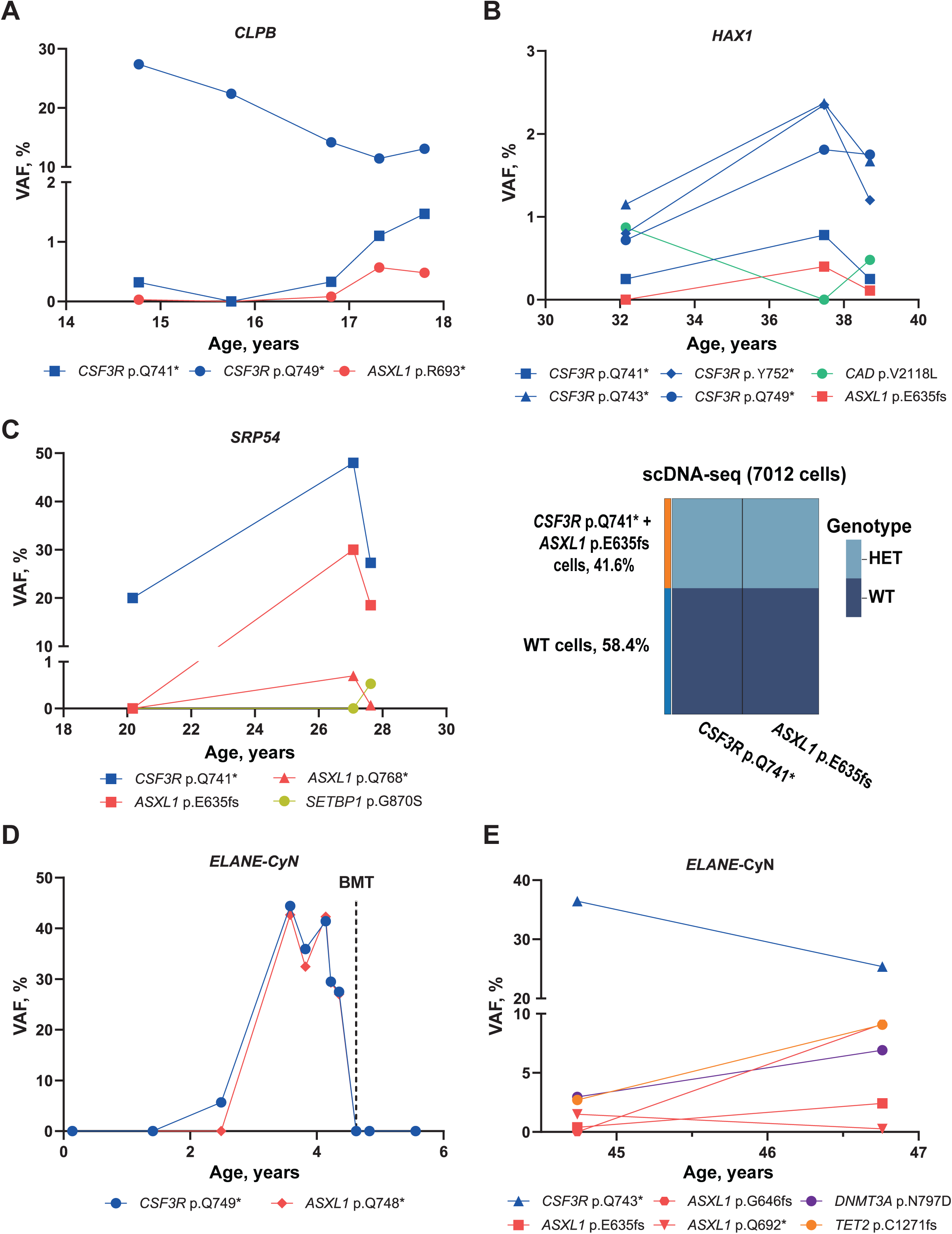
Clonal dynamics and single-cell architecture of somatic clones in leukemia-free CN patients, predominantly with *CSF3R* and *ASXL1* mutations. **A-B,** Longitudinal monitoring of somatic mutations in *CLPB* and *HAX1* patients. **C,** Time-course analysis of VAF for somatic clones (left) and in scDNA-seq of 7012 cells (right) in a *SRP54* patient, showing the co-occurrence of *CSF3R* p.Q741* and *ASXL1* p.E635fs mutations within the same clone. The height of each colored segment represents the proportion of cells assigned to a specific mutant (HET) or wild-type (WT) genotype. **D-E**, Time-course analysis of VAF for somatic clones in two *ELANE*-CyN patients. In a first *ELANE*-CyN patient (**D**), a dashed line marks time point of bone marrow transplantation (BMT). In a second *ELANE*-CyN patient (**E**) additional age-related mutations (*TET2*, *DNMT3A*) emerged alongside *CSF3R* and *ASXL1* alterations. Abbreviations: ANC, absolute neutrophil count; VAF, variant allele frequency; scDNA-seq, single-cell DNA sequencing; HET, heterozygous; WT, wild-type.

*TP53*-driven CH was detected in 50% of leukemia-free SDS patients, consistent with previously reported rates of 45-48%^10,24^ (**Figure 3A**). The p.R175H hotspot mutation^27^ was the most frequent variant, detected in seven SDS patients (**Figure 3B**). *TP53* VAFs were significantly lower in leukemia-free SDS patients than in samples collected before progression to MDS/AML (median interval to diagnosis, 1.33 years; Mann-Whitney U test, *p =* 0.0002) (**Figure 3C**). In our exploratory receiver operating characteristic (ROC) analysis, a VAF threshold of 0.074 for the dominant *TP53* clone discriminated SDS prior to MDS/AML from leukemia-free SDS (AUC = 0.99, 95% CI: 0.95–1.00, *p* = 0.00013) with 96.3% specificity, 100% sensitivity, 80.0% positive predictive value (PPV), and 100% negative predictive value (NPV). Individual SDS patients carried up to eight distinct *TP53* mutations (**Figure S2A**). In addition to *TP53*, SDS patients carried a range of low-prevalence somatic mutations, including *DNMT3A, IDH1*, *PTPN11* and *RUNX1*. Notably, no *CSF3R* or *ASXL1* mutations were detected (**Figure 3A**).

**Figure 3.**
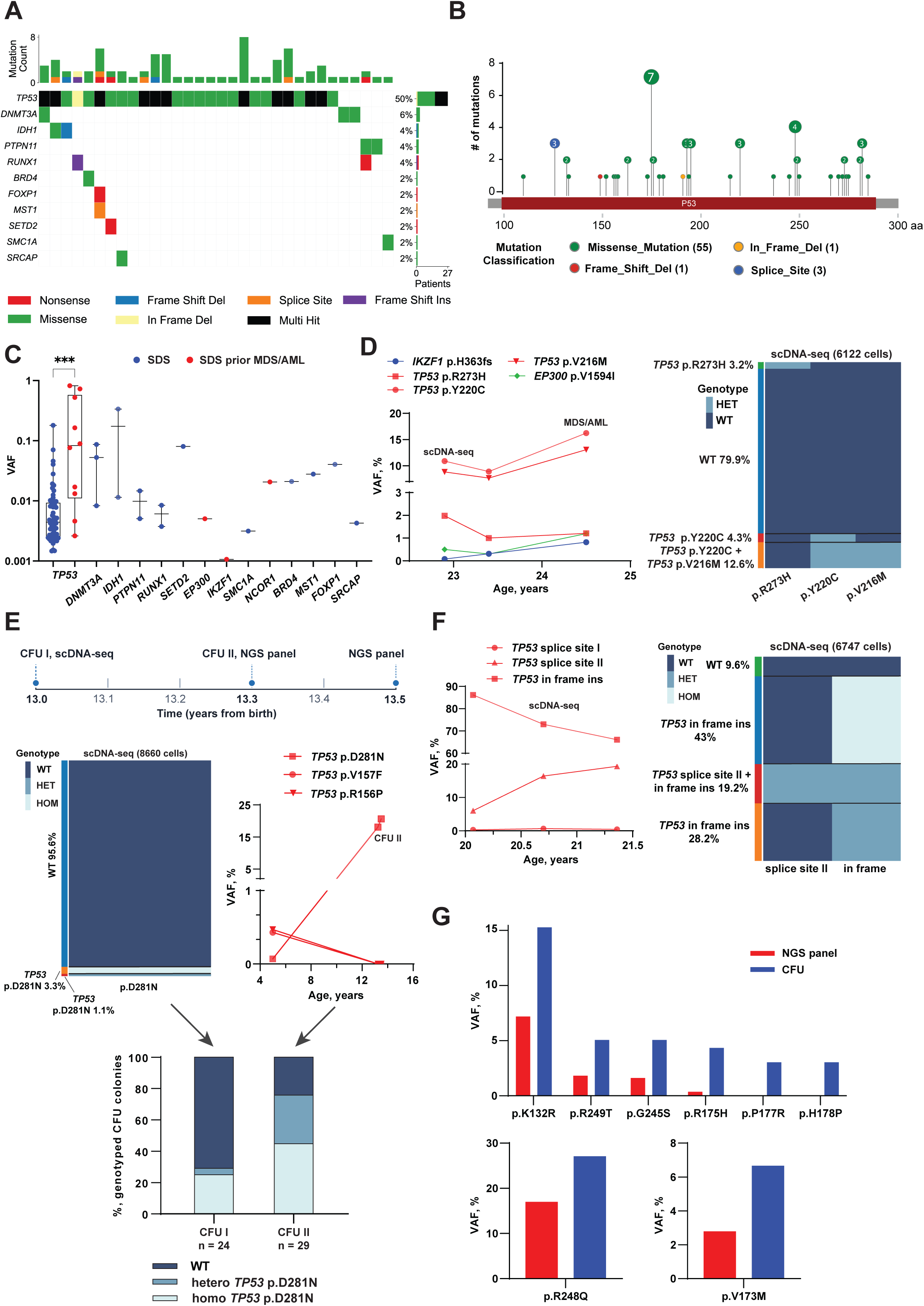
CH and biallelic *TP53* mutations in SDS patients. **A**, Oncoplot of somatic mutations in SDS patients at leukemia-free time points (n = 54), highlighting the predominance of *TP53* alterations. Top bar plots indicate individual mutation counts per SDS patient. Each column represents a patient with SDS. Mutation types are color-coded according to the legend key. Only patients with somatic mutations are shown. **B**, Lollipop plot with frequencies of *TP53* mutations detected in leukemia-free SDS patients. All identified *TP53* mutations are localized within the *TP53* DNA-binding domain. Mutations are color-coded by type. Numbers within the circles indicate the number of distinct *TP53* mutations at each residue. The p.R175H hot-spot mutation was most frequently detected (n = 7). **C**, Comparison of somatic VAFs between leukemia-free *SDS* patients (dark blue) and patients prior to MDS/AML (red). Statistical comparison was restricted to *TP53*. *TP53* VAFs significantly differed between the SDS and SDS prior MDS/AML groups (Mann-Whitney test, *p =* 0.0002). The y-axis is log10 scaled to resolve low-frequency variants. Each point is one somatic mutation, plotted at its median VAF across the patient’s samples. **D**, VAF tracking in an SDS patient showing the stable presence and expansion of *TP53* clones (p.V216M, p.Y220C, and p.R273H) (left). Right: corresponding scDNA-seq revealed that the p.V216M and p.Y220C mutations co-occurred in a compound heterozygous state within the same clone (12.6%) 1.5 years prior to malignant transformation. The height of each colored segment represents the proportion of cells with a given mutant (HET) or wild-type (WT) genotype. **E,** Clonal evolution of the *TP53* p.D281N clone in a SDS patient, assessed by scDNA-seq, panel sequencing, and CFU assay. Upper part: schematic representation of a timeline indicating when specific assays were performed. Middle right part: time-course analysis of somatic *TP53* mutations measured by NGS panel. Only the *TP53* p.D281N clone exhibited selective expansion over time, with significantly increasing frequency. Middle left part: single-cell heatmap of 8660 cells analyzed by scDNA-seq. The height of each colored segment represents the proportion of cells with a given mutant (HOM or HET) or WT genotype. scDNA-seq identified hetero- and homozygous *TP53* p.D281N clones. Lower part: validation of scDNA-seq and NGS panel results through Sanger sequencing of CFU colonies. The bar charts show the proportions of WT (dark blue), heterozygous (blue), and homozygous (light blue) *TP53* p.D281N CFU colonies. Between the first and second observations, the proportion of homozygous colonies increased from 25% to 44.8%, while heterozygous colonies expanded from 4.17% to 31.0%. **F,** VAF tracking (left) and scDNA-seq (right) of a third SDS patient identified complex clonal architecture at the single-cell level, namely founding heterozygous cells and the emergence of a large homozygous *TP53* clone (43%) and a compound heterozygous clone (19.2%) harboring a second splice-site mutation. The height of each colored segment represents the proportion of cells with a given mutant (HOM or HET) or WT genotype. **G,** Comparison of *TP53* VAF between the NGS panel (red) and genotyping of CFU colonies (dark blue) of three SDS patients. CFU assay (upper part) identified heterozygous *TP53* mutations p.P177R and p.H178P that were missed by the NGS panel. Abbreviations: SDS, Shwachman-Diamond syndrome; VAF, variant allele frequency; HOM, homozygous; HET, heterozygous; CFU, colony-forming unit; NGS, next generation sequencing.

Progression to MDS/AML in SDS is strongly associated with acquisition of biallelic *TP53* alterations^24^. Using scDNA-seq in three SDS patients with high *TP53* VAF, we detected biallelic *TP53* clones months to years before malignant transformation (**Figure 3D-F**). In the first SDS patient, panel sequencing revealed three *TP53*, one *IKZF1*, and one *EP300* mutations. scDNA-seq showed that two of the TP53 variants, p.V216M and p.Y220C, were present in compound heterozygous form 1.5 years before leukemogenic transformation (**Figure 3D**). Of three *TP53* mutations initially detected in the second SDS patient, only the p.D281N clone expanded over eight years, from 0.07% to 20.6% VAF (**Figure 3E**). scDNA-seq of CD34⁺ cells identified 3.3% heterozygous and 1.1% homozygous p.D281N cells. Genotyping of individual CFU colonies confirmed these findings, tracking a rapid three-month expansion, from 25% to 44.8% for homozygous and from 4.17% to 31.0% for heterozygous clones (**Figure 3E**). In the third SDS patient with biallelic *TP53* mutations, panel sequencing identified a dominant clone carrying a *TP53* in-frame insertion and the emergence of a second clone with a splice-site *TP53* mutation, but only scDNA-seq resolved the underlying architecture: founding heterozygous cells, a large homozygous *TP53* clone comprising 43% of cells, and a compound heterozygous clone (19.2%) carrying an additional splice-site mutation (**Figure 3F**).

We next compared CFU-based single-cell genotyping with time-matched panel sequencing in three SDS patients (**Figure 3G**). CFU genotyping consistently yielded higher *TP53* mutation frequencies than panel sequencing. In one patient, CFU genotyping additionally identified two heterozygous *TP53* mutations (p.P177R and p.H178P) that were not detected by panel sequencing. These results indicate that single-cell genotyping enables sensitive monitoring of *TP53* clonal evolution in SDS.

### Clinical determinants, detection across sample sources, and age dynamics of CH in CN and SDS

We previously reported that CN patients requiring high G-CSF doses (>8 µg/kg/day) and with an ANC below 2.19× 10^9^/L are at increased risk of malignant transformation ^22^. In the present study, CH rates in *ELANE* and *HAX1* patients -the two largest genetic CN subgroups in our cohort - did not increase with G-CSF dose, at a threshold of 10 µg/kg/day, or 20 µg/kg/day (**Figure 4A**, **S2B**). Neither the maximal dose before sampling nor the maximal lifetime G-CSF dose was associated with CH. Likewise, ANC was not associated with CH or *CSF3R* mutations in *ELANE* and *HAX1* patients (**Figure 4B, S2C**), or with CH and *TP53* mutations in the SDS cohort (**Figure 4C, S2D**).

**Figure 4.**
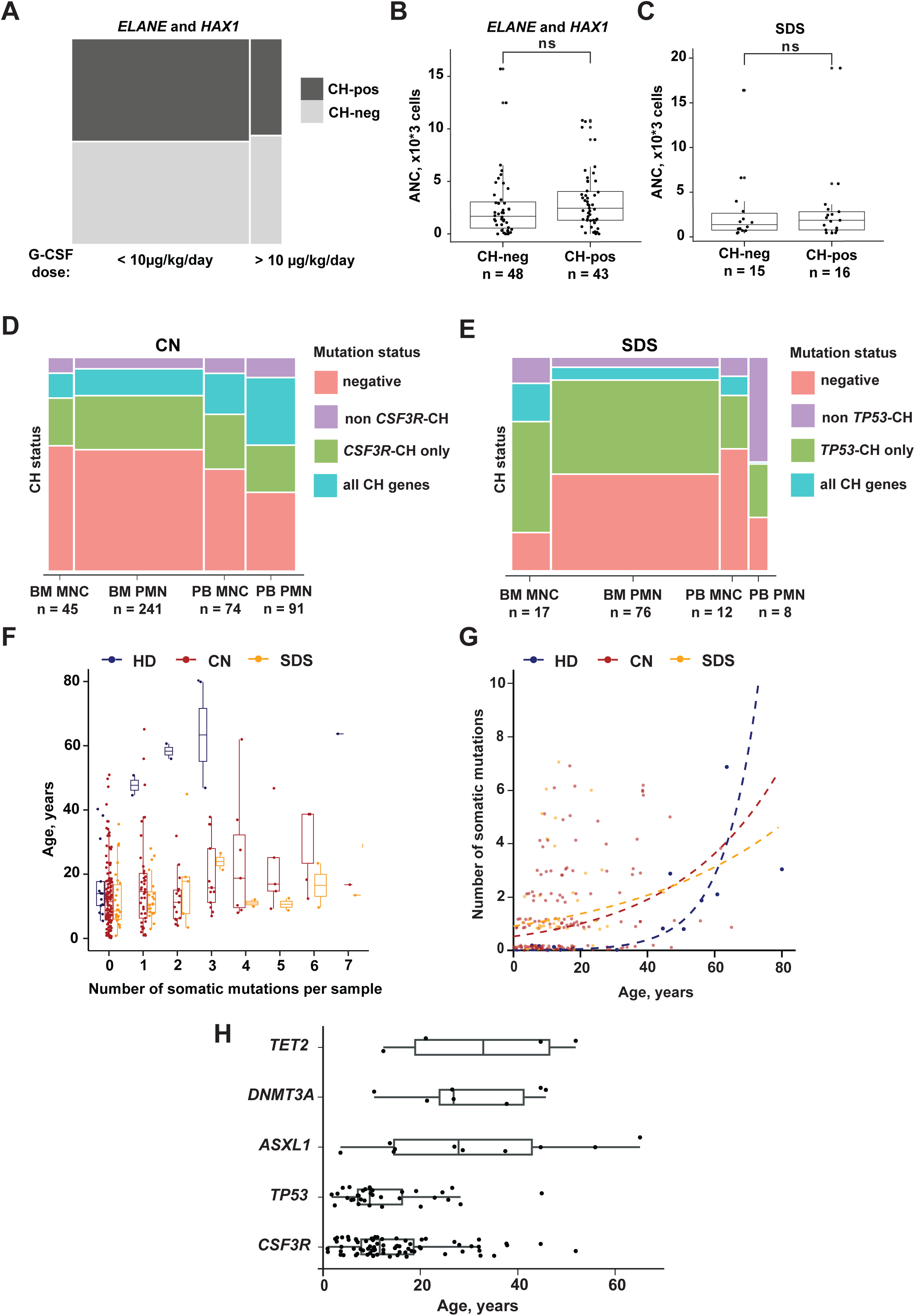
Correlation of CH with clinical parameters, cell sources, and age of CN and SDS patients. **A,** CH prevalence and G-CSF dose. Mosaic plot showing the distribution of CH-positive (dark grey) and CH-negative (light grey) cases in *ELANE* and *HAX1* cohorts stratified by daily dose of G-CSF (10 µg/kg/day cutoff). No significant increase in CH rates was observed in a group with G-CSF doses over 10 µg/kg/day (FET, OR 0.89, *p* = 1). **B,** Correlation between absolute neutrophil counts (ANCs) and CH status. ANCs are plotted for CH-negative and CH-positive *ELANE* and *HAX1* patients. The difference in ANCs was not statistically significant. **C,** ANC and CH status in SDS patients. ANC levels are plotted for CH-negative and CH-positive patients within the SDS cohort. No significant association was found between CH and ANC. **D,** Mosaic plot shows the proportion of CH-positive samples across different sample materials (BM vs. PB) and cell fractions (MNC vs. PMN) in the CN cohort. **E,** Mosaic plot shows the proportion of CH-positive samples across BM and PB fractions in SDS patients. BM MNCs showed the highest proportion of CH-positive cases. **F,** Boxplot showing the number of acquired mutations (x-axis) versus age (y-axis), grouped by healthy (blue), CN (red) and SDS (orange) individuals, clearly demonstrates CH at an early age in CN and SDS patients. **G**, Chart of same data as in panel F, with mutation count (y-axis) plotted against age (x-axis). Dashed lines display the modelled mutations counts for healthy (blue), CN (red) and SDS (orange) individuals through the range of ages. **H**, Box plots illustrate the age at detection for somatic mutations across five CH-related genes in CN and SDS patients. Individual data points represent independent mutational events. Abbreviations: ANC, absolute neutrophil count; BM, bone marrow; PB, peripheral blood; MNC, mononuclear cells; PMN, polymorphonuclear cells; G-CSF, granulocyte colony-stimulating factor.

To assess the feasibility of blood-based CH monitoring, we compared the somatic VAFs and the proportion of CH-positive cases between MNC and PMN fractions from BM and PB. In CN patients, median VAFs were highest in BM MNC (**Figure S2E**), but modelled VAFs did not suggest a significant shift in detection power between BM and PB (odds ratio 1.19, 95% CI: 0.82-1.73, p-value 0.351) nor between MNC and PMN fractions (odds ratio 1.20, 95% CI: 0.79-1.87, p-value 0.396). CH-positive cases were likewise not enriched in BM (**Figure 4D**). In paired BM and PB samples, VAFs of *CSF3R* and other key drivers in blood samples, especially in PB PMN closely mirrored those in BM (**Figure S3**). PB is therefore suitable for routine CH monitoring in CN patients. In SDS, VAF did not differ significantly between PB and BM or PMN and MNC fractions for any CH-related gene (predominantly *TP53*) (**Figure S4A**). The proportion of CH-positive cases was highest in BM MNC, but the difference between BM vs PB was not statistically significant (p-value 0.8) (**Figure 4E**). In the one SDS patient with paired BM and PB samples, the dominant *TP53* clones had lower VAFs in blood, but all *TP53* mutations detected in BM were also found in blood (**Figure S4B**).

CH is rare in healthy individuals below 40 years of age and increases in frequency thereafter^28,29^. In CN and SDS patients, in contrast, CH was already present in childhood and adolescence (**Figure 4F**). We therefore modeled the dynamics of somatic mutation acquisition in patients and healthy donors. In the healthy cohort, the expected mutation count started at 0.008 at birth and increased by a factor of 1.10 per year (95% CI: 1.04 -1.21), reaching one mutation per individual by age 50. In contrast, in CN and SDS patients, the modelled intercept at birth was higher (0.53 and 0.91, respectively), whereas the annual rate of mutation acquisition was lower than in healthy donors (rate ratio 0.94, 95% CI: 0.86-0.99, *p*-value 0.029 for CN; 0.93, 95% CI: 0.84-0.99, *p*-value 0.032 for SDS) (**Figure 4G**). In line with the low mutation burden observed in MDS/AML samples, these data suggest that CH in CN and SDS patients is driven by selective expansion of HSPCs carrying specific mutations -most frequently *CSF3R* in CN and *TP53* in SDS - rather than by an elevated mutation rate. *CSF3R* and *TP53* mutations were acquired predominantly within the first two decades of life, whereas mutations in age-related CH genes (*TET2*, *DNMT3A*) appeared later (**Figure 4H**).

### Somatic mutation composition and transformation risk in CN patients preceding MDS/AML

We next analyzed samples prior to leukemia, available for 28 out of 37 patients who developed MDS/AML. For comparison, we selected samples from leukemia-free patients with the same germline background, including *ELANE, HAX1, WAS, JAGN1, SLC37A4,* and *SBDS* (**Table S1**, **Figure 5A-B**). The median collection time prior to overt MDS/AML was 4.08 years. *CSF3R* and *RUNX1* mutations were enriched in prior to leukemia samples. The odds ratio (OR) for *CSF3R* was 5.1 (95% CI 1.7 - 16.8; *p*-value 8.6 x 10^-4^) whereas no *RUNX1* mutations were observed in the analyzed CN leukemia-free cohort (**Figure 5C**). *ASXL1* mutations were more frequent, but the difference was not significant (OR 4.3, 95% CI 0.6 - 24.2, p-value 0.074) (**Figure 5C**). A significant difference in clone size between the groups was observed for *CSF3R* mutations only (**Figure 5D**). By integrating results of our previous *CSF3R* cDNA sequencing^9^ with current data, we evaluated the impact of *CSF3R* mutation status on the progression from CN stage to MDS/AML. *CSF3R* mutations significantly increased the risk of leukemic transformation (log-rank test chisq 3.98, *p*-value 0.046) (**Figure 5E**). In CN patients, each additional mutation was associated with an increased risk of progression to MDS/AML (OR 1.40, 95% CI 1.10 - 1.75, *p*-value 0.0028). The association was even stronger when the number of mutated genes, rather than individual mutations, was taken into account (OR 2.6, 95% CI 1.7 - 4.26, *p*-value 3 x 10^-5^) (**Figure 5F**).

**Figure 5.**
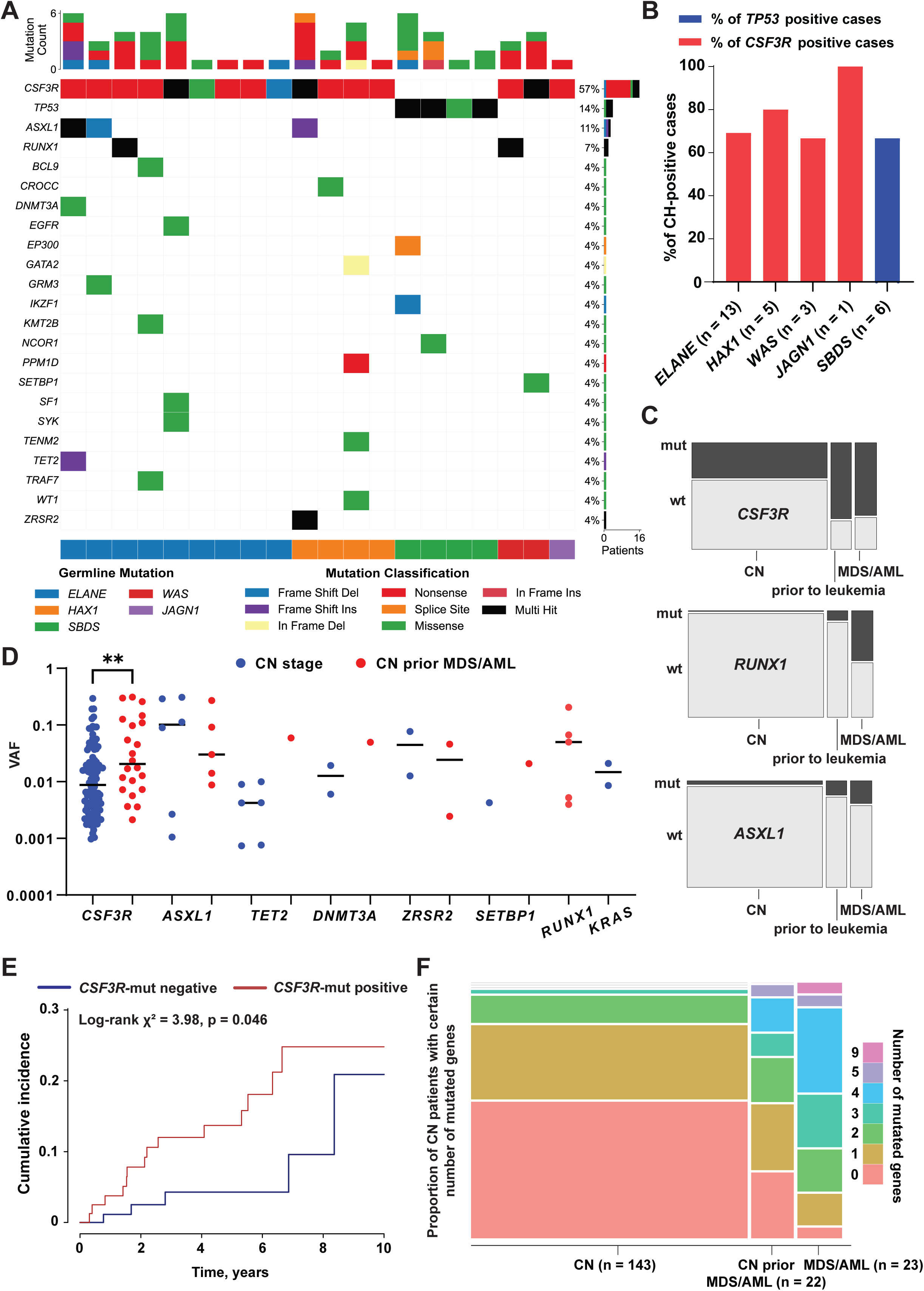
Genetic predictors of leukemogenic transformation in CN and SDS patients. **A,** Somatic mutational landscape of CN-MDS/AML and SDS-MDS/AML patient samples prior to leukemic transformation. Top bar plots indicate individual mutation counts per patient. Each column represents a patient. Neutropenia types and mutation classifications are color-coded according to the legend key. Only patients with somatic mutations are shown. **B,** CH prevalence prior to leukemic transformation across genetic groups. The bar chart illustrates the percentage of *CSF3R*-positive (red) and *TP53*-positive (dark blue) cases, stratified by germline background, in samples collected prior to the onset of leukemia. **C,** Enrichment of key mutations in the samples prior to the onset of leukemia. The proportions of *CSF3R*, *RUNX1*, and *ASXL1* mutated cases in leukemia-free CN patients (CN, n = 143), in samples prior to leukemia from CN-MDS/AML patients (n = 22) and at MDS/AML stage (n = 23) are shown in dark grey. *CSF3R* (FET, OR 5.1, 95% CI 1.7 - 16.8; *p =* 8.6 x 10^-4^) and *RUNX1* (FET, OR ∞, 95% CI 1.3 - ∞; *p =* 0.017) mutations were significantly enriched in samples prior to leukemia collected from CN-MDS/AML patients. The prevalence of *ASXL1* mutations did not differ significantly between the leukemia-free CN and samples prior to the onset of leukemia (FET, OR = 4.3, 95% CI 0.6 - 24.2; *p* = 0.074). **D,** Comparison of VAFs for leukemia driver genes between patients at the CN stage (dark blue) and the prior to leukemia samples (red). Statistical comparison was restricted to *CSF3R* and *ASXL1*. A statistically significant increase in median clone size was observed only for *CSF3R* mutations (Mann-Whitney test, \*\**adjusted p* < 0.01). Each point is one somatic mutation, plotted at its median VAF across the patient’s samples. **E,** Impact of *CSF3R* status (80 positive, 101 negative cases) on the risk of progression to AML/MDS: Kaplan-Meier curves showing the cumulative MDS/AML incidence stratified by the presence (red) or absence (dark blue) of *CSF3R* mutations. Among the analyzed CN cohort, 19 patients developed MDS/AML. Presence of *CSF3R* mutations is associated with an increased risk of leukemic transformation (Log-rank test, chisq = 3.98; p = 0.046). **F,** Number of mutated genes and progression risk. Mosaic plot shows the proportion of CN patients with varying numbers of mutated genes across three stages: CN, CN prior to MDS/AML, and overt MDS/AML. Each additionally mutated gene significantly increases the odds of progression to MDS/AML by 2.6 (95% CI 1.7 - 4.26; *p =* 3 x 10^-5^). Abbreviations: FET, Fisher’s exact test; VAF, variant allele frequency; SDS, Shwachman-Diamond syndrome; CN, congenital neutropenia.

### Clonal and cytogenetic evolution to MDS/AML in CN

We next determined the spectrum of somatic mutations in CN-MDS/AML patients. *CSF3R* mutations were present in the majority of cases (63%), followed by *RUNX1* (41%), *ASXL1* (22%), *PTPN11* (15%), and *SETBP1* (15%) (**Figure 6A**), confirming the previously reported cooperativity between *CSF3R* and *RUNX1* mutations^12^. The somatic mutation spectrum did not differ between germline phenotypes, with the exception of SDS, in which *TP53* mutations predominated. *RUNX1, PTPN11,* and *SETBP1* mutations were present in leukemia samples but rarely detected during the CH phase, indicating that these genes are acquired late in leukemogenic transformation. The most common genetic pattern in CN-MDS/AML is the co-occurrence of *CSF3R* and *RUNX1* mutations, with or without additional *ASXL1* mutations. Consistent with a hierarchical relationship, eight CN-MDS/AML patients without acquired *CSF3R* mutations carried no *RUNX1*, *SETBP1* and *ASXL1* mutations, except for one patient with a missense *RUNX1* mutation. These findings indicate that *CSF3R* mutations provide the clonal background on which *RUNX1* and *ASXL1* mutations expand. In a representative *ELANE*-CyN case, the leukemic clones at diagnosis carried *CSF3R* p.Q754*, *ASXL1* p.S955fs, *RUNX1* p.R139fs, with a minor tetrasomy 21 subclone (**Figure 6B, S5A**). At relapse, scDNA-seq and panel sequencing revealed three subclones: one with tetrasomy 21 affecting the mutant *RUNX1* allele, and two carrying distinct *KRAS* mutations (p.G60V and p.G12V) (**Figure 6B**). Interestingly, analysis of sporadic MDS/AML cases (n=91) with nonsense mutations in *CSF3R* showed that, even without a germline background, truncating *CSF3R* mutations preferentially cooperate with mutated *ASXL1* and *RUNX1* (**Figure S5B**).

**Figure 6.**
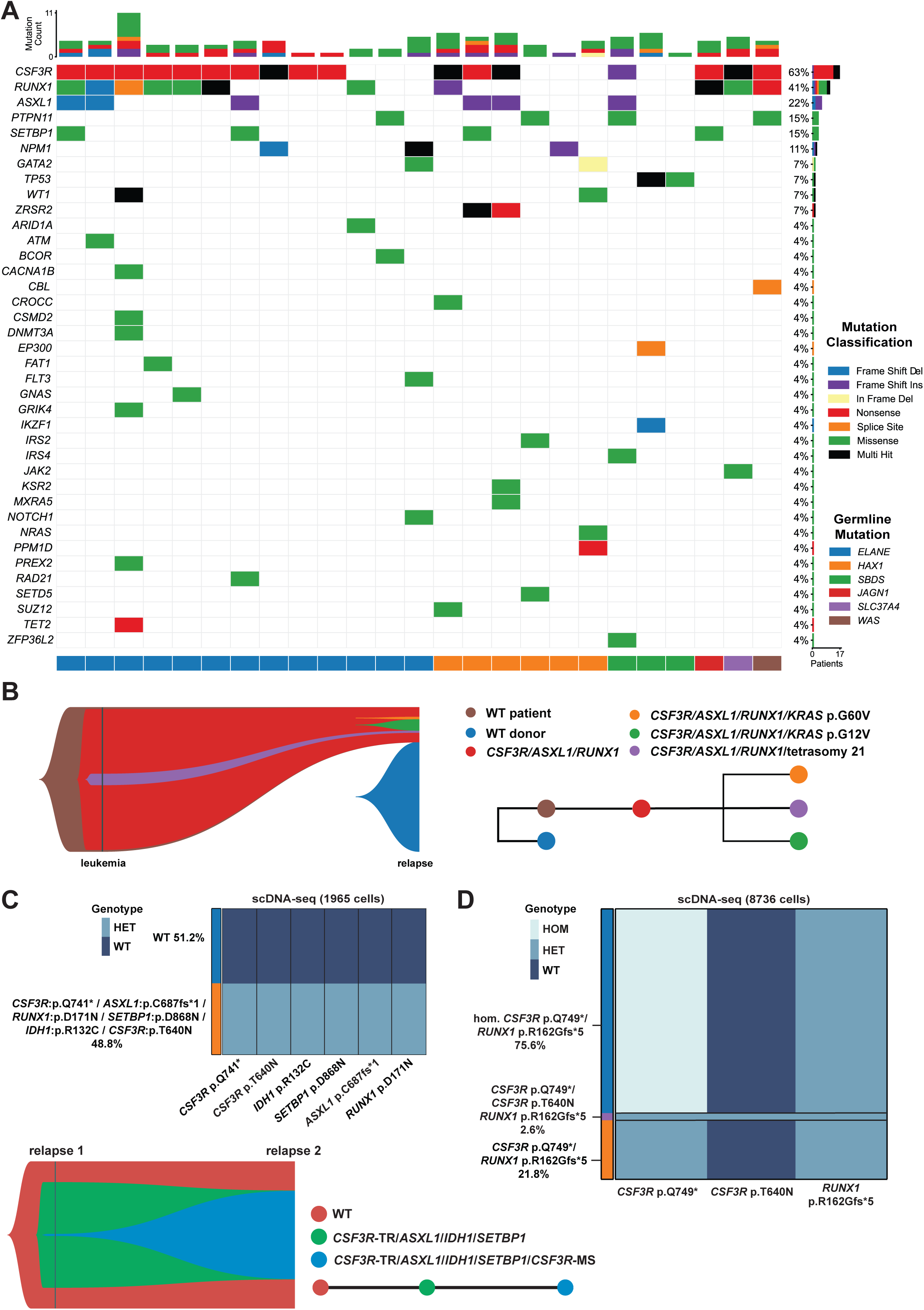
Somatic mutation landscape and clonal evolution in CN-MDS/AML and SDS-MDS/AML patients. **A,** The spectrum of acquired mutations in MDS/AML samples from CN and SDS patients (n = 27). Top bar plots indicate individual mutation counts per patient. Each column represents a patient. Underlying germline mutations are indicated by the color track. Mutation types are color-coded according to the legend key. Only patients with somatic mutations are shown. **B,** Clonal evolution (fish plot) of a CyN patient *ELANE* p.V101M with leukemia relapse assessed by scDNA-seq. Relapse is characterized by evolution of the main leukemic clone with *CSF3R* p.Q754\**, ASXL1* p.S955fs, and *RUNX1* p.R139fs mutations, with subclones co-harboring *KRAS* p.G12V (green, dominant subclone), *KRAS* p.G60V (orange, minor subclone), or tetrasomy 21 (purple, minor subclone). **C,** Upper part: scDNA-seq of a CyN-AML patient *ELANE* p.A233P, p.V235Wfs*1 at relapse revealed that 48.8% of cells belong to a dominant clone harboring six mutations: *CSF3R* p.Q741* and p.T640N, *ASXL1* p.C687fs*1, *RUNX1* p.D171N, *SETBP1* p.D868N, and *IDH1* p.R132C. Lower part: fish plot of clonal evolution between the first and second relapse samples analyzed by scDNA-seq. *RUNX1* p.D171N is not displayed in the fish plot because it was not covered by the scDNA-seq analysis of the first relapse sample. **D,** Clonal architecture of a CN/AML patient *HAX1* p.W44* with missense and biallelic stop-codon mutations in *CSF3R*. A single cell heatmap of 8736 cells shows the presence of the largest clone (75.6%) with homozygous *CSF3R* p.Q749* and heterozygous *RUNX1* p.R162Gfs*5 mutations. A smaller clone (2.6%) has acquired an additional activating *CSF3R* mutation (p.T640N), while a third clone (21.8%) remains heterozygous for the founding mutations. Abbreviations: CyN, cyclic neutropenia; CN, congenital neutropenia; scDNA-seq, single-cell DNA sequencing; TR, truncating mutation; MS: missense mutation.

In three CN patients with leukemia, we detected membrane-proximal *CSF3R* mutations (p.T615A, p.T618I, p.T640N) alongside nonsense *CSF3R* mutations. These proximal mutations were not detected during the CH stage. Longitudinal tracking combined with scDNA-seq resolved this sequence in two patients. In an *ELANE*-CyN patient, leukemia onset was marked by the emergence of a *RUNX1-* and *SETBP1-*mutant clone on a pre-existing background of truncating *CSF3R* and *ASXL1* mutations (**Figure S1D**). At relapse, the clone had acquired *CSF3R* p.T640N, *IDH1* p.R132C, and *BCOR* p.R810*, together with numerous low-frequency mutations. In the earliest relapse sample, the *CSF3R* p.T640N mutation was detected at a VAF of 0.7% (**Figure S1D**). scDNA-seq showed that the main leukemic clone comprised 70.7% of cells, with the *CSF3R* double-mutant subclone present in 1% (**Figure S6A**). Eleven months later, this *CSF3R* p.Q741*/p.T640N clone had expanded to 48.8% of cells (**Figure 6C**). In a second *HAX1*-MDS/AML patient, scDNA-seq identified three clones: a dominant clone with homozygous *CSF3R* p.Q749* and *RUNX1* p.R162Gfs*5 mutations in 75.6% of cells; a second clone with heterozygous *CSF3R* p.Q749* and *RUNX1* p.R162Gfs*5 mutations in 21.8% of cells; and a minor subclone that had additionally acquired *CSF3R* p.T640N mutation in 2.6% of cells (**Figure 6D**).

We also evaluated the prevalence of two recurrent chromosomal aberrations in 30 out of 37 SDS- and CN-MDS/AML patients, for whom karyotype data were available: monosomy 7 or del(7q) (-7/del(7q)) and trisomy 21 (**Table S6, Figure S6B**). -7/del(7q) status was available for all 30 patients and trisomy 21 status for 28 patients. Trisomy 21 was detected in 10 patients (35.7%) and -7/del(7q) in 15 patients (50.0%), with 5 patients harboring both abnormalities. Of the remaining patients who lacked both of these aberrations, 4 exhibited additional cytogenetic findings, and 6 had a normal karyotype. *CSF3R* and *RUNX1* mutations were identified in leukemia samples irrespective of cytogenetic status. *ASXL1* mutations did not co-occur with isolated monosomy 7, and *PTPN11* mutations were found predominantly in cases with trisomy 21 (**Figure S6C**).

Analysis of nucleotide substitutions across patient groups revealed an enrichment of G>A transitions in CN, reflecting the prevalence of stop-gain mutations at glutamine residues within *CSF3R* hotspots (**Figure 1C**, **S6D**). In SDS, C>T and T>C transitions predominated in *TP53* (**Figure S6D**). In the COSMIC database, C>T transitions are the second most frequent substitution type in *TP53* after G>A.

## Discussion

Here we present a comprehensive analysis of CH in the largest cohort of patients with congenital neutropenia syndromes studied to date, spanning CN, CyN and SDS. CH was highly prevalent, affecting 37.7% of CN and 60% of SDS patients. Consistent with previous reports^9,30,31^, somatic *CSF3R* mutations predominated in CN, whereas *TP53* mutations were most frequent in SDS. CyN patients acquired a similar mutation spectrum to CN patients, but at lower prevalence, in line with their lower leukemia incidence and likely reflecting differences in disease pathophysiology. While *CSF3R* and *TP53* mutations were mutually exclusive across most genetic subgroups, they co-occurred in patients with *SRP54*- or *CLPB*-associated CN, suggesting that the germline defects determine which clones are selected.

Importantly, CH prevalence did not differ between patients receiving high versus low G-CSF doses and was not correlated with ANC, indicating that G-CSF dose does not drive the emergence of CH, which is consistent with previously published results^9,10,31^. We previously reported, however, that the hazard of MDS/AML increases 1.24-fold (95% CI: 1.08-1.43-fold) per doubling of the G-CSF dose in CN patients ^22^. Since the endpoint in that analysis was leukemic transformation rather than CH, higher G-CSF requirements may influence transformation of existing clones rather than their initial emergence. Alternatively, a high G-CSF requirement may itself be a marker of more severe defect in HSPCs, with transformation risk determined by the degree of HSPCs stress. Both possibilities warrant further investigation. Survival analysis of time to leukemic transformation demonstrated that the presence of a *CSF3R* mutation is associated with an increased risk of leukemia in CN patients. However, *CSF3R-*mutant clones can persist for many years without progression to leukemia^12,30^. *ASXL1* mutations emerged much earlier in CN and CyN patients than in individuals with age-related CH, and the overall mutation burden remained largely stable over decades. These observations suggest that transformation to MDS/AML in CN and SDS is driven primarily by positive selection of clones carrying specific drivers, likely under the intrinsic cellular stress caused by the germline defects, rather than by gradual mutation accumulation typical for aging.

Among leukemia-free CN patients, *ASXL1* was the second most frequently mutated gene after *CSF3R*, followed by much rarer *TET2* and *DNMT3A* mutations. In 7 out of 10 cases, *ASXL1* mutations emerged on a *CSF3R* mutant background, potentially establishing a sub-clonal population that drives leukemogenesis. *TP53*, *SETBP1* and *RUNX1* mutations were found in only isolated cases in the leukemia-free CN cohort. Longitudinal analysis showed that *ASXL1* mutations were detectable years before leukemogenic transformation, whereas *RUNX1* and *SETBP1* mutations were acquired shortly before disease progression, suggesting that they represent late events that drive rapid transformation to MDS/AML.

In CN-MDS/AML patients, the most frequently mutated genes were *CSF3R, RUNX1*, *ASXL1*, *PTPN11* and *SETBP1*. These mutations were typically acquired on a *CSF3R*-mutant background and often co-occurred in combinations, suggesting the emergence of highly transformative subclones. *PTPN11* mutations were acquired independently of *CSF3R* in two out of four cases. scDNA-seq of relapsed patients revealed that the dominant leukemic clones at relapse derived from the *CSF3R*-mutant backbone already established before transformation. The co-occurrence of *CSF3R* with *ASXL1*, *RUNX1* and *SETBP1* therefore defines a high-risk molecular profile, and its detection – even at low VAF – should lead to intensified surveillance and early consideration of allogeneic HSCT.

The practical value of CH monitoring depends on whether it can be performed on blood rather than bone marrow. In CN, the dominant *CSF3R* clones were detected in both BM and PB in all paired samples. In SDS, the trend towards higher *TP53* VAFs in BM MNC suggests that blood may be less sensitive for small *TP53* clones, which is relevant given the low VAF threshold for progression risk. For other genes, the limited sample size precludes definitive conclusions.

Understanding why *RUNX1* and *ASXL1* mutations frequently co-occur with *CSF3R* mutations during leukemogenic transformation is of particular importance. *ASXL1* mutations cooperate with activating *CSF3R* mutations (T618I/T621I) to create a pre-leukemic chronic neutrophilic leukemia (CNL)-like state^32^, associated with aggressive disease and poorer survival^33^. Mechanistically, the epigenetic and transcriptional dysregulation triggered by *ASXL1* or *RUNX1* mutations may accelerate myeloid transformation by amplifying the transcriptional changes driven by mutant *CSF3R*. Consistent with this, sporadic leukemia cases harboring *CSF3R* truncating mutations most frequently co-occur with *ASXL1* or *RUNX*1 mutations.

We identified membrane-proximal activating *CSF3R* mutations in combination with truncating mutations at the CN/AML stage, but not in samples collected prior to leukemia. Unlike truncating variants, which hyperactivate G-CSF-dependent signaling, membrane-proximal mutations cause ligand-independent receptor activation^34,35^. Clones carrying both mutation types would therefore combine constitutive receptor activation with loss of signal termination. This "super-driver" state may contribute to the rapid clonal expansion observed in one patient, from 1% to 48.8% of cells within eleven months. Homozygous truncating CSF3R mutations, detected in one patient, may have a comparable effect.

We defined a *TP53* VAF threshold of > 0.074 for predicting progression to MDS/AML in SDS, enabling monitoring of malignant transformation using NGS panels. In patients who transformed, biallelic *TP53* mutant clones were detectable in SDS patients before MDS/AML, consistent with findings reported by Shimamura’s group.^24^ The interval between detection of biallelic mutations and overt disease ranged from months to years, meaning that detection identifies risk without indicating how soon transformation will occur. We demonstrated that *TP53* zygosity can be assessed by Sanger sequencing of single-cell-derived CFU colonies, which requires no specialized platform and is therefore available to any laboratory performing CFU assays. Notably, this approach detected *TP53* mutations missed by panel sequencing. This indicates that bulk VAF does not fully capture clonal composition, particularly for small or subclonal *TP53* mutations.

The limitations of this study include the following: despite being the largest cohort reported to date, the CN/AML group remains small, which limits the precision of mutation frequency estimates; sampling intervals were not standardized, which may bias the detection of rapidly evolving clones in less frequently monitored patients; the *TP53* VAF threshold for predicting progression to MDS/AML was derived and tested in the same cohort and requires independent validation before clinical use; we also did not investigate complex structural variants.

In summary, leukemogenic transformation in CN, CyN and SDS arises from early selection of specific clones that persist for years and acquire late drivers shortly before overt MDS/AML. The mutations marking this transformation, *CSF3R*, *ASXL1* with *RUNX1* and/or *SETBP1* in a majority of CN and CyN, or biallelic *TP53* in SDS, are detectable in peripheral blood before transformation, providing a basis for risk-adapted surveillance and timely consideration of allogeneic HSCT.

## Acknowledgment

We would like to thank the patients, their parents, and the physicians from the Severe Chronic Neutropenia International Registry (SCNIR) and the European Working Group of Myelodysplastic Syndromes (EWOG-MDS) networks for their cooperation in providing the patients’ material for this study. This work was supported by the Madeleine Schickedanz-Kinderkrebsstiftung, the de.NBI Cloud within the German Network for Bioinformatics Infrastructure (de.NBI) and ELIXIR-DE, the German Federal Ministry of Education and Research (BMBF), the German Cancer Aid (Deutsche Krebshilfe), the German Jose Carreras Leukemia Foundation, and the National Insitutes of Health (NIH). We thank Ms. K. Hähnel and Ms. U. Hermanutz-Klein for excellent technical support, Mr. J. Herold and Ms. M. Otto for help with Sanger sequencing genotyping of CFU colonies, Mr. D. Frank for his contribution to summarization of patient genetic data, and Dr. Patricia Arreba-Tutusaus for her assistance with the preparation of the manuscript and figures, and Dr. Sergey Mastitsky for consulting on the statistics. Prof. Akiko Shimamura is gratefully acknowledged for her valuable comments and review of the manuscript.

## Author Contributions

M.K., J.S. and S.K. made initial observations and designed the experiments; M.K., and S.K. analyzed the data; M.K., J.S., K.W., S.K., C.Z and S. M.-H. interpreted the results; K.W., C.Z., S.M.-H., Y.L.B., D.C.D., V.M., M.Ke., A.A.B., C.M.N., M.E., A.Y.-N. M.M., P.L. and C.L. provided samples from patients and healthy donors, as well as clinical, genetic, and cytogenetic data. M.H. collected and summarized data on ANC, G-CSF dose, and cytogenetics. T.H. and N.N. provided data on sporadic leukemia cases with *CSF3R* mutation. M.K., J.S., S.K., and K.W. wrote the manuscript. J.S. acquired funding, provided resources, and supervised the project. All authors reviewed and approved the final version of the manuscript.

