## Supplementary_methods_and_legends for "Deciphering the Dynamics of Leukemogenic Transformation from Clonal Hematopoiesis in Congenital Neutropenia"

**Supplementary information**

**Material and methods**

**DNA isolation**

Bone marrow mononuclear cells (BM MNCs), bone marrow polymorphonuclear cells (BM PMNs), peripheral blood mononuclear cells (PB MNCs), and peripheral blood polymorphonuclear cells (PB PMNs) were isolated using Ficoll-Hypaque Plus gradient centrifugation (GE Healthcare, Cat. No. 17-1440-03). $CD34^+$ cells were separated from BM MNCs through positive selection using the EasySep™ Human CD34 Positive Selection Kit II (STEMCELL Technologies, Cat. No. 17856), following the standard protocol. $CD34^+$ cells with a purity of over 80%, as determined by flow cytometry, were used in the study. The QIAamp DNA Blood Mini (Qiagen, Cat. No. 51104) or Midi (Qiagen, Cat. No. 51183) kits were used for DNA purification.

**Library preparation**

DNA samples were sonicated to achieve fragments of approximately 150-300 bp using Covaris M220 focused-ultrasonicator (PerkinElmer). End repair and adaptor ligation were performed using xGen™ Prism DNA Library Prep Kit (IDT, Cat. No. 10006202, 10006203) and xGen™ cfDNA & FFPE DNA Library Prep v2 MC (IDT, Cat. No. 10010206, 10010207) followed by purification with AMPure XP beads (Beckman Coulter, Cat. No. A63880). Fixed single-stranded unique molecular identifiers (UMIs) sequences were added to the insert during the first ligation step. Sample index barcodes (xGen UDI Primers Plate, Cat. No. 10005922) were introduced via library amplification by PCR. Quality and quantity assessments of sequencing libraries were performed using a Qubit fluorometer (Thermo Fisher Scientific) and the Agilent 2100 Bioanalyzer system (Agilent Technologies). Next, DNA libraries were quantified using NEBNext® Library Quant Kit for Illumina (New England Biolabs, Cat. No. E7630S).

Target enrichment was performed according to the manufacturer's protocol for xGen® Hybridization and Wash Kit (IDT, Cat. No. 1080577) or xGen® Hybridization and Wash v2 Reagents (IDT, Cat. No. 10010351) together with xGen® Hybridization and Wash v2 Beads (IDT, Cat. No. 10010353). Briefly, 19 to 23 sample DNA libraries per sequencing run were equimolarly pooled and hybridized with biotin-labeled xGen® Lockdown Probes targeting specific genomic regions, with human Cot DNA and xGen® Universal Blockers TS (IDT, Cat. Nr. 1075474), added to minimize off-target binding. Streptavidin beads were used to capture hybridized DNA, followed by sequential washes to remove nonspecifically bound fragments. Post-capture PCR was performed to amplify the enriched library, and AMPure XP® beads (Beckman Coulter, Cat. No. A63880) were applied for final purification. Pooled enriched libraries were quantified and qualified using Qubit fluorometer (Thermo Fisher Scientific) and the Agilent 2100 Bioanalyzer system (Agilent Technologies) to ensure proper concentration and fragment size for Illumina sequencing.

**Sequencing of sample libraries**

Sample libraries were sequenced in paired-end 150 mode on the Illumina NovaSeq platform. On average, 12 Gb of raw data was generated per sample after enrichment with capture panel version 1 and 9 Gb of raw data per library after enrichment with capture panel version 2. We aimed to reach >1000x duplex depth of coverage and variants mutation detection sensitivity of 0.1% allele frequency or less.

**Raw reads preprocessing**

Raw sequenced reads were converted to unaligned BAM files using Picard (http://broadinstitute.github.io/picard/v2.27.1) FastqToSam. The UMI part of the sequenced reads was extracted into the RX tag using fgbio (https://github.com/fulcrumgenomics/fgbio) v2.0.1 ExtractUmisFromBam. After UMI extraction, reads were converted back to FASTQ format with Picard SamToFastq and mapped to hg38 reference with bwa[1] v0.7.17. The resulting alignments were merged back to unmapped reads with Picard MergeBamAlignment to preserve tags and read group information.

We additionally used sequences of the UMIs provided by the manufacturer to correct errors in mapped reads' RX tags using fgbio CorrectUmis with the following parameters: --max-mismatches=3 and --min-distance=1.

Reads with the same UMI tag and mapped location were grouped using fgbio GroupReadsByUmi with parameters --strategy=paired, --edits=0 and --min-map-q=20. Reads originating from complementary strands of the primary DNA molecule were algorithmically collapsed into highly accurate duplex consensus reads utilizing fgbio CallDuplexConsensusReads with the parameters --error-rate-pre-umi=45, --error-rate-post-umi=30, and --min-input-base-quality=30. Duplex consensus reads were remapped to the hg38 reference genome and merged with unmapped reads to retain tags and read group information.

**Variant calling and generation of the Panel of Normals**

We first called variants in 20 cord blood donors (24 samples) with GATK[2] and Mutect2[3] v4.2.6.1 to capture recurrent technical artifacts and formed a Panel of Normals (PoN) with GATK CreateSomaticPanelOfNormals. It was then used for variant calling in patient samples.

**Variant calling in patient samples**

To adjust sensitivity to the unusually high read depth and the small number of reads supporting the true variant allele in both PoN and patients' samples, we used the following non-default Mutect2[3] parameters: --dont-use-soft-clipped-bases true, --max-mnp-distance 0, --max-reads-per-alignmentstart 0, --mitochondria-mode true, --recover-all-dangling-branches true, --pcr-indel-model NONE, --annotations-to-exclude AS_StrandBiasMutectAnnotation and --af-of-alleles-not-in-resource 0. We further filtered raw Mutect2 calls with GATK FilterMutectCalls, increasing --max-events-in-region to 5 to retain true variants occurring in different clones of the same sample at mutational hotspots. All low frequency somatic mutations (1-2 patients per mutation) were manually curated using the Integrative Genomics Viewer (IGV)[4].

**Variant calling in sequential patient samples**

In cases where multiple consequent samples from the same donor were sequenced, we additionally increased the sensitivity of the variant calling for known true mutations in that donor. To do so, we combined the filtered variants calls from all samples of that donor using bcftools[5] v1.15.1 merge. The resulting vcf file was used as a set of alleles to force-call in all samples from the donor, regardless of supporting evidence. Such forced variant calls were not further filtered and were annotated directly. By doing so, we were able to retrospectively re-confirm the clone at the earliest stages of its expansion or during its decline.

**Variants annotation**

Detected variants were annotated using GATK Funcotator against full funcotator annotation data source v1.7. We additionally annotated each variant with its number of occurrences in the following cBioPortal[6] leukemic cohorts: aml_ohsu_2018, mds_tokyo_2011, mds_iwg_2022, laml_tcga_pan_can_atlas_2018, mds_mskcc_2020, laml_tcga, aml_target_2018_pub, laml_tcga_pub. While point mutations required an exact coordinate match to consider a match with public data, a 5 bp proximity window was permitted for indels to account for minor alignment shifts.

**Exploratory analysis and visualization of variants**

Functionally annotated somatic candidates in MAF format were loaded into R v4.2.2 (R Core Team, 2024. R: A Language and Environment for Statistical Computing. R Foundation for Statistical Computing, Vienna, Austria. (https://www.R-project.org)) for analysis and visualization. Variants with gnomAD[7]v2.1 populational frequency exceeding 0.0001 were filtered out except ones with Likely Pathogenic, Pathogenic and Pathogenic/Likely Pathogenic ClinVar[8] v20180429_hg38 annotations (**Table S5)**. We further removed germline variant calls that passed Mutect2 filters by using variant allele frequency (VAF) threshold 0.26. Such a threshold was not applied for MDS/AML, post-BMT, and relapse samples, since true somatic clones could have expanded. The VAF filter was not applied to the next category of mutations: annotated as Likely Pathogenic, Pathogenic and Pathogenic/Likely Pathogenic by ClinVar; mutations with overlapping COSMIC[9] v85 records (Primary histology = haematopoietic_neoplasm); nonsense mutations and frameshift/inframe indels. Manual curation of both kept and filtered out mutations resulted in lifting of filters for some patients or blacklisting of genes/loci from further analysis (available upon request).

Data exploratory analysis was performed using R packages *dplyr* (https://CRAN.R-project.org/package=dplyr) v1.4.1, *tidyr* (https://CRAN.R-project.org/package=tidyr) v1.3.1, (https://CRAN.R-project.org/package=purrr) v1.0.2 and visualized with *ggplot2* (https://CRAN.R-project.org/package=ggplot2) v3.5.1. and *ggmosaic* (https://CRAN.R-project.org/package=ggmosaic) v0.3.3. For plots with necessary genomic contexts such as lollipop plot and oncoplot – we used g3viz[10] v 1.2.0, *maftools*[11] v2.22.10 and *oncoplotpy* (https://github.com/kandabarau/oncoplotpy) v1.0.0.

**Statistical modelling**

Effect of age and disease on clonal hematopoiesis was tested with the Negative Binomial Generalized Linear Model with R package MASS (https://CRAN.R-project.org/package=MASS) v7.3-60.2. Model efficiency was assessed and effects visualized with *interactions* (https://CRAN.R-project.org/package=interactions) v1.1.5 and *DHARMa* (https://CRAN.R-project.org/package=DHARMa) v0.4.6 packages. The distribution of mutations, mutated genes, and patient positivity rates across discrete phenotypic subgroups, as well as the sensitivity and variance of CH detection across diverse sample materials was tested by fitting Generalized Linear Models (GLM) with stats R-base package. Overrepresentation of patients with mutation in a specific gene in progressive group compared to non-progressive was tested using Fisher's exact test. Group differences in VAF were tested with the two-sided Mann-Whitney U test. ROC curves and AUC analysis were computed with GraphPad Prism 11.1.0.

**Single cell DNA sequencing (scDNA-seq)**

scDNA-seq library preparation was performed using the Mission Bio Tapestri platform V2, according to the Tapestri Single-Cell DNA Sequencing User Guide (PN 3354). The Myeloid Panel (https://www.missionbio.com/products/panels/myeloid) was utilized to analyze mutations. The panel encompasses 45 genes with 312 amplicons, covering mutations relevant to diverse types of leukemia. BM MNC, PB MNC or $CD34^+$ cells with viability exceeding 80% were prepared at a concentration of 3000–4000 cells/µL. Sequencing was performed on the Illumina NovaSeq platform using PE150/PE250 sequencing with at least 70 Gb of raw data per sample.

**scDNA-seq data processing and variant calling**

Somatic variants detected by capture panel version 1 and version 2 were lifted over to hg19 using the *GenomicRanges*[12] v1.55.4 and *liftOver* v1.27.0 packages for single-cell level analysis (https://www.bioconductor.org/help/workflows/liftOver). scDNA-seq FASTQ files were uploaded to https://eu.tapestripipeline.missionbio.com/ and processed with Tapestri Pipeline DNA v2.0.2 (Mission

Bio, Inc.). Mutations and cells were filtered, and clones were further decomposed with Mission Bio mosaic command line tools v3.1.1, employing hg19 known sample mutations as a whitelist.

**Progression-free survival analysis**

We analyzed progression-free survival (PFS) intervals for patients with and without *CSF3R*-mutated clonal hematopoiesis (CH). For the *CSF3R*-wildtype cohort, the index date was defined as the date of the first sample collection; for the *CSF3R*-CH cohort, the index date was the date of the first *CSF3R*-positive sample. For patients who progressed to MDS/leukemia, the event date was defined as the first date of MDS or leukemia diagnosed. For the remaining cases, the censoring date was determined by the date of last follow-up, the date of bone marrow transplantation (BMT), or the study end date, whichever occurred first. A 10-year cutoff was applied to maintain data density and mitigate the late-stage data sparsity. Differences in progression-free survival distributions were evaluated with a log-rank test of *survival* base R package.

**Colony-forming unit (CFU) assay and Sanger sequencing of CFU colonies**

To generate single cell-derived colonies from $CD34^+$ cells, CFU assays were performed using a semi-solid methylcellulose matrix. $CD34^+$ cells from SDS patients were purified from bone marrow (BM) via magnetic bead selection and frozen. After thawing, $CD34^+$ cells were resuspended in Iscove's Modified Dulbecco's Medium (IMDM) supplemented with 2% Fetal Bovine Serum (FBS) and plated at a concentration of $1 \times 10^3$ to $2 \times 10^3$ cells per 35 mm dish in enriched MethoCult medium (Stemcell Technologies, Cat. No. H4435) containing rhSCF, rhGM-CSF, rhIL-3, rhIL-6, rhG-CSF, and rhEPO.

The cells were incubated at 37°C in a humidified $CO_2$ incubator for 14 days. Individual colonies were classified based on their morphological characteristics. For molecular analysis, individual colonies, primarily CFU-granulocyte (CFU-G), CFU-monocyte (CFU-M), and CFU-granulocyte-macrophage (CFU-GM) colonies, were isolated from the methylcellulose matrix. Approximately 20-35 CFU colonies were picked per SDS patient. Isolated cell colonies were lysed in 35 µl of Quick extract DNA solution (Lucigen, Cat. No. 20802) and incubated 15 min at 65°C and 15 min at 98°C. The *TP53* mutation status of each colony was subsequently determined via Sanger sequencing. Sequencing primers used for this analysis are available upon request.

**Supplementary figures legends**

**Supplementary figure 1. CH across genetic subgroups and its dynamics in CN patients.** **A,** Frequency of CH with VAF > 2% across CN genetic subgroups, SDS patients and genetically unclassified cases (neg_exome). Bar charts represent the percentage of patients in each genetic category that tested positive for CH with cutoff VAF >2%. **B,** Distribution of somatic *CSF3R* mutation counts per CN patient. **C,** Time-course analysis of VAF for somatic clones in a *SRP54* patient, showing the presence of *CSF3R* p.Q754* and *TP53* p.R282L mutations. **D**, Clonal evolution and leukemic transformation in a CyN patient with *ELANE* p.A233P and p.V235Wfs*1. VAF heatmap for patient samples spanning 8.4 years is shown. The heatmap tracks the progression of a *CSF3R* and *ASXL1* mutant clone from the neutropenia stage to leukemia and relapse, highlighting the late-stage acquisition of the activating *CSF3R* p.T640N mutation and numerous low-frequency subclonal mutations. **E,** Heatmap of VAFs detected in samples from a patient with *HAX1* p.W44* over 5.6 years prior to malignant transformation. The data demonstrates the early presence of a founding *CSF3R* p.Y752* clone and the subsequent emergence of *ASXL1* and *ZRSR2* mutations 2.3 and 1.3 years before overt leukemia, respectively. **F,** Persistent CH driven by *ASXL1* mutations was tracked in a *JAGN1* patient for 6.75 years without progression to leukemia. Abbreviations: VAF, variant allele frequency; CH, clonal hematopoiesis; CN, congenital neutropenia; AML, acute myeloid leukemia.

**Supplementary figure 2. *TP53* mutation counts in SDS, CH and G-CSF dose, ANC, and VAF in BM and PB cell fractions in CN and SDS patients.** **A,** Distribution of *TP53* mutation counts in SDS patients, showing the number of distinct somatic *TP53* mutations per SDS patient. **B,** CH prevalence, depending on therapeutic G-CSF dose. Mosaic plot showing the distribution of CH-positive (dark grey) and CH-negative (light grey) cases in *ELANE* and *HAX1* CN cohorts stratified by daily dose of

G-CSF (20 μg/kg/day cutoff). No significant increase in CH rates was observed in a group with G-CSF dose over 20 μg/kg/day (FET, OR = 1.02, $p$ = 1.0). **C,** ANC and *CSF3R* mutation status. ANCs are plotted for *CSF3R*-negative and *CSF3R*-positive *ELANE*-CN and *HAX1*-CN patients. The difference in ANCs was not statistically significant. **D,** Correlation between ANC and *TP53* mutation status in SDS patients. ANC levels are plotted for *TP53*-negative and *TP53*-positive patients within the SDS cohort. No significant association was found between the presence of *TP53* mutation and ANC. **E**, Comparison of VAF across BM and PB fractions (MNC and PMN) for the CN cohort. Median VAF values for each fraction are indicated above the boxplots. Total number (n) of mutations detected is added to the x axis labels. Abbreviations: ANC, absolute neutrophil count; BM, bone marrow; PB, peripheral blood; MNC, mononuclear cells; PMN, polymorphonuclear cells; VAF, variant allele frequency; CH, clonal hematopoiesis; CN, congenital neutropenia; SDS, Shwachman-Diamond syndrome.

**Supplementary figure 3. Comparison of somatic mutation burden in BM and PB cell fractions in paired samples from *HAX1*-CN (A, B), *ELANE*-CN (C), and *ELANE*-CyN (D) patients.** The bar plots show somatic mutation burdens (measured as % of VAF) across four distinct cell fractions: peripheral blood mononuclear cells (PB MNC), peripheral blood polymorphonuclear cells (PB PMN), bone marrow mononuclear cells (BM MNC), and bone marrow polymorphonuclear cells (BM PMN). Bar colors indicate specific somatic mutations, as defined in the legends below each panel.

**Supplementary figure 4. *TP53* VAF in BM and PB cell fractions from SDS patients. Comparison of somatic mutation burden in BM and PB cell fractions in paired samples from SDS patients.** **A**, Comparison of VAF across BM and PB fractions (MNC and PMN) for the SDS cohort. Median VAF values for each fraction are indicated above the boxplots. Total number (n) of mutations detected is added to the x axis labels. **B,** The bar plots show somatic mutation burdens (measured as % VAF) across four distinct cell fractions: PB MNC, PB PMN, BM MNC, and BM PMN. Bar colors indicate specific somatic mutations, as defined in the legends below each panel. Abbreviations: BM, bone marrow; PB, peripheral blood; MNC, mononuclear cells; PMN, polymorphonuclear cells; VAF, variant allele frequency; SDS, Shwachman-Diamond syndrome.

**Supplementary figure 5. Clonal evolution and leukemic transformation in a CyN patient and spectrum of somatic mutations in sporadic leukemia cases with truncating mutations in *CSF3R*. A,** Heatmap showing VAFs of somatic and germline variants across eight sequential samples from a CyN patient with *ELANE* p.V101M who developed leukemia and relapse. Samples are ordered chronologically from 6.4 years of pre-diagnosis to 2.4 years of post-diagnosis. Variants are color-coded: patient germline (yellow), *CSF3R* frameshift (white), initiating leukemic (orange), donor-specific (light blue), and relapse-acquired mutations (red). **B,** Mutational spectrum of sporadic *CSF3R*-mutant MDS/AML. Oncoplot shows the somatic mutations in a cohort of 91 sporadic cases with truncating *CSF3R* mutations. Individual patient mutation counts are provided in the top bar plot. Specific mutation types are denoted by color-coded boxes.

**Supplementary figure 6. Clonal architecture of CyN-AML patient sample that carries both *CSF3R* p.Q741* and p.T640N mutations. Monosomy 7 and trisomy 21 in SDS- and CN-MDS/AML patients. Patterns of single nucleotide substitutions in CN and SDS patients. A,** scDNA-seq analysis of 501 cells from a CyN-AML patient with *ELANE* p.A233P and p.V235Wfs*1 at clinical relapse. Segment height indicates the relative proportion of cells assigned to each genotype. A dominant clone (70.7%) harbored four mutations: *CSF3R* p.Q741*, *ASXL1* p.C687fs*1, *IDH1* p.R132C, and *SETBP1* p.D868N. A minor subclone (1%) additionally acquired missense mutation *CSF3R* p.T640N. The *RUNX1* p. D171N mutation detected in this patient by panel sequencing was not covered by the current scDNA-seq analysis. **B**, The proportion of monosomy 7 or del(7q) (-7/del(7q), purple) and trisomy 21 (+21, green) in 37 SDS- and CN-MDS/AML patients. N/A: karyotype data is not available. **C,** Horizontal stacked bar chart shows the distribution of cytogenetic subgroups (normal karyotype, trisomy21 (+21), monosomy 7/del(7q) (-7/del(7q)), concurrent (+21 and -7), or other chromosomal abnormalities, across somatic mutations detected in SDS- and CN-MDS/AML patients. Genes are ranked by total mutation frequency. The cytogenetic status of each mutated case is color-coded. **D,** Comparison of single nucleotide substitution profiles for the predominant CH drivers (*CSF3R* for CN; *TP53* for SDS) and all other mutated genes (aggregated) in CN and SDS patient samples.
