## Supplementary_figures for "Deciphering the Dynamics of Leukemogenic Transformation from Clonal Hematopoiesis in Congenital Neutropenia"

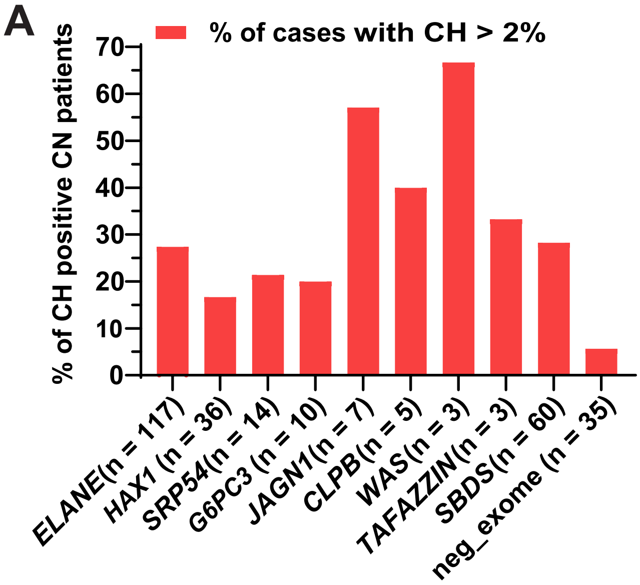


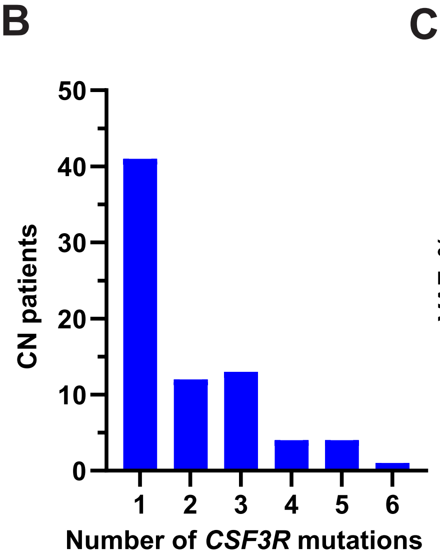


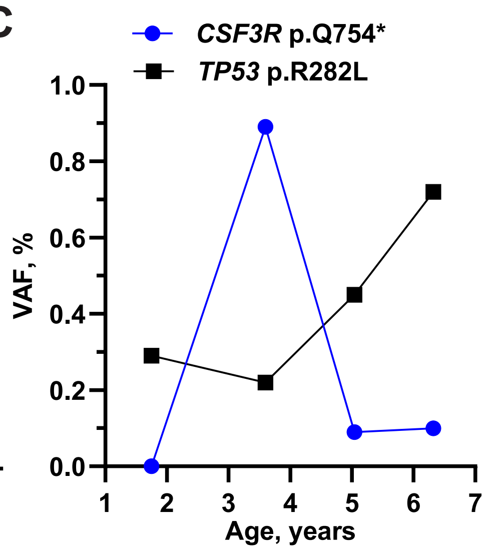


D

VAF: 0 – 0,6

| Gene | Mutation | Neutropenia stage | | Leukemia | Leukemia – Relapse | |
|---|---|---|---|---|---|---|
| | | sample 1 | sample 2 | sample 3 | sample 4 | sample 5 |
| *CSF3R* | Q741* | 0.070 | 0.039 | 0.181 | 0.424 | 0.369 |
| *ASXL1* | C687fs | 0 | 0.03 | 0.157 | 0.428 | 0.366 |
| *RUNX1* | D171N | 0 | 0 | 0.098 | 0.595 | 0.500 |
| *SETBP1* | D868N | 0 | 0 | 0.182 | 0.444 | 0.395 |
| *IDH1* | R132C | 0 | 0 | 0 | 0.434 | 0.390 |
| *GRIK4* | G546G | 0 | 0 | 0 | 0.376 | 0.405 |
| *CSF3R* | T640N | 0 | 0 | 0 | 0.007 | 0.378 |
| *SETBP1* | V231L | 0 | 0 | 0 | 0.049 | 0.102 |
| *GRIK2* | M867I | 0 | 0 | 0 | 0 | 0.101 |
| *CSMD1* | T2038S | 0 | 0 | 0 | 0 | 0.100 |
| *IDH2* | intronic | 0 | 0 | 0 | 0 | 0.099 |
| *CROCC* | intronic | 0 | 0 | 0 | 0 | 0.047 |
| *CSMD1* | A2052V | 0 | 0 | 0 | 0 | 0.038 |
| *MAP2* | M269I | 0 | 0 | 0 | 0 | 0.026 |
| *SUZ12* | E594* | 0 | 0 | 0 | 0 | 0.020 |
| *TP53* | C141F | 0 | 0 | 0 | 0 | 0.019 |
| *GRM3* | R612Q | 0 | 0 | 0 | 0 | 0.018 |
| *PKD1L2* | intronic | 0 | 0 | 0 | 0 | 0.007 |
| *PRDM16* | P633L | 0 | 0 | 0 | 0.041 | 0 |
| *BCL9* | intronic | 0 | 0 | 0 | 0.060 | 0 |
| *TET2* | S268S | 0 | 0 | 0 | 0.044 | 0 |
| *FLT3* | T227M | 0 | 0 | 0 | 0.092 | 0 |
| *BCOR* | R810* | 0 | 0 | 0 | 0.423 | 0 |
| *BCORL1* | P503P | 0 | 0 | 0 | 0.032 | 0 |
| **Months from leukemia** | | **-65.3** | **-39.2** | **0** | **25** | **35.8** |

E

VAF: 0 – 0.5

| Gene | Mutation | Neutropenia stage | | | Leukemia |
|---|---|---|---|---|---|
| | | sample 1 | sample 2 | sample 3 | sample 4 |
| *CSF3R* | Y752* | 0.0026 | 0.34 | 0.36 | 0.26 |
| *ASXL1* | G646fs | 0 | 0.29 | 0.3 | 0.26 |
| *ZRSR2* | Splice_Site | 0 | 0 | 0.045 | 0.43 |
| *CSF3R* | Q743* | 0.02 | 0.023 | 0.0066 | 0 |
| *CSF3R* | Q754* | 0 | 0.013 | 0 | 0 |
| *ZRSR2* | W111* | 0 | 0 | 0 | 0.095 |
| *FRYL* | A2513A | 0 | 0 | 0 | 0.03 |
| *SETBP1* | G870S | 0 | 0 | 0 | 0.0074 |
| Months from leukemia | | **-67.3** | **-27.5** | **-15.4** | **0** |

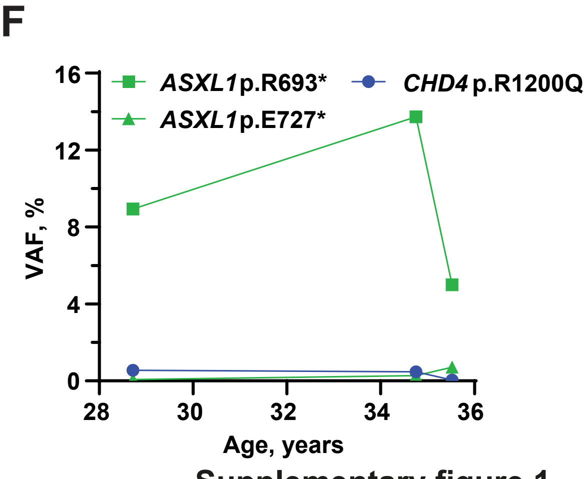


Supplementary figure 1

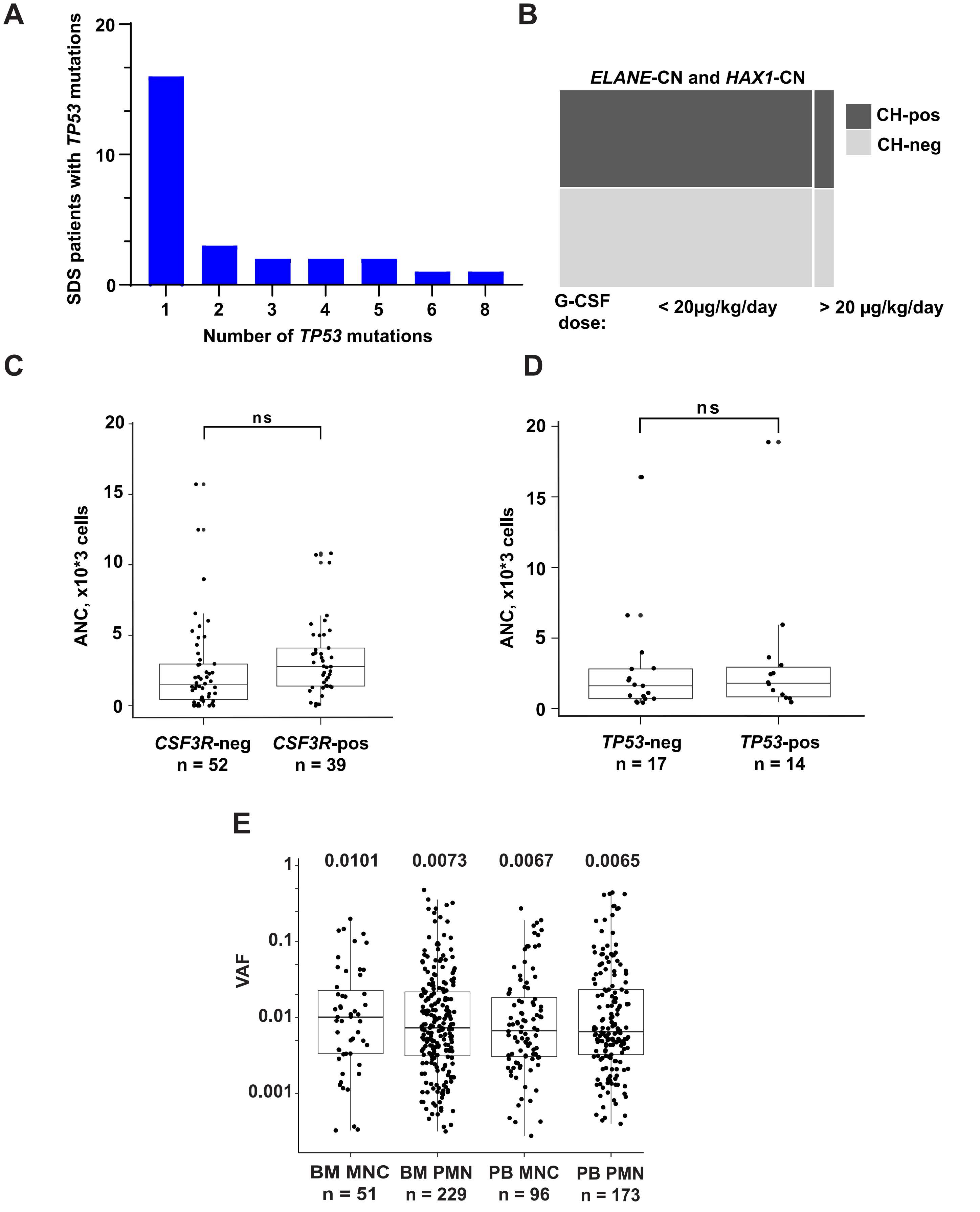


Supplementary figure 2

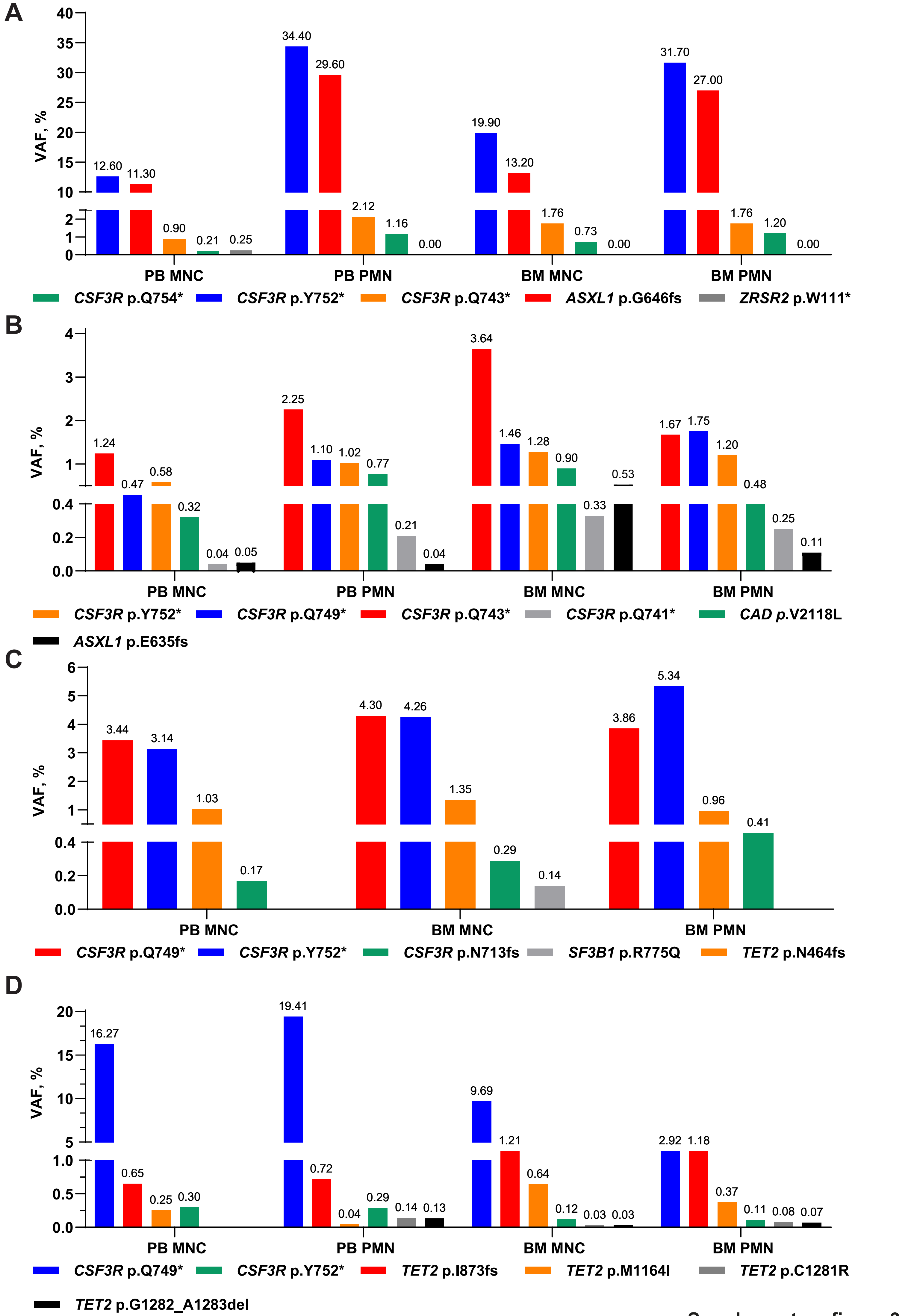


Supplementary figure 3

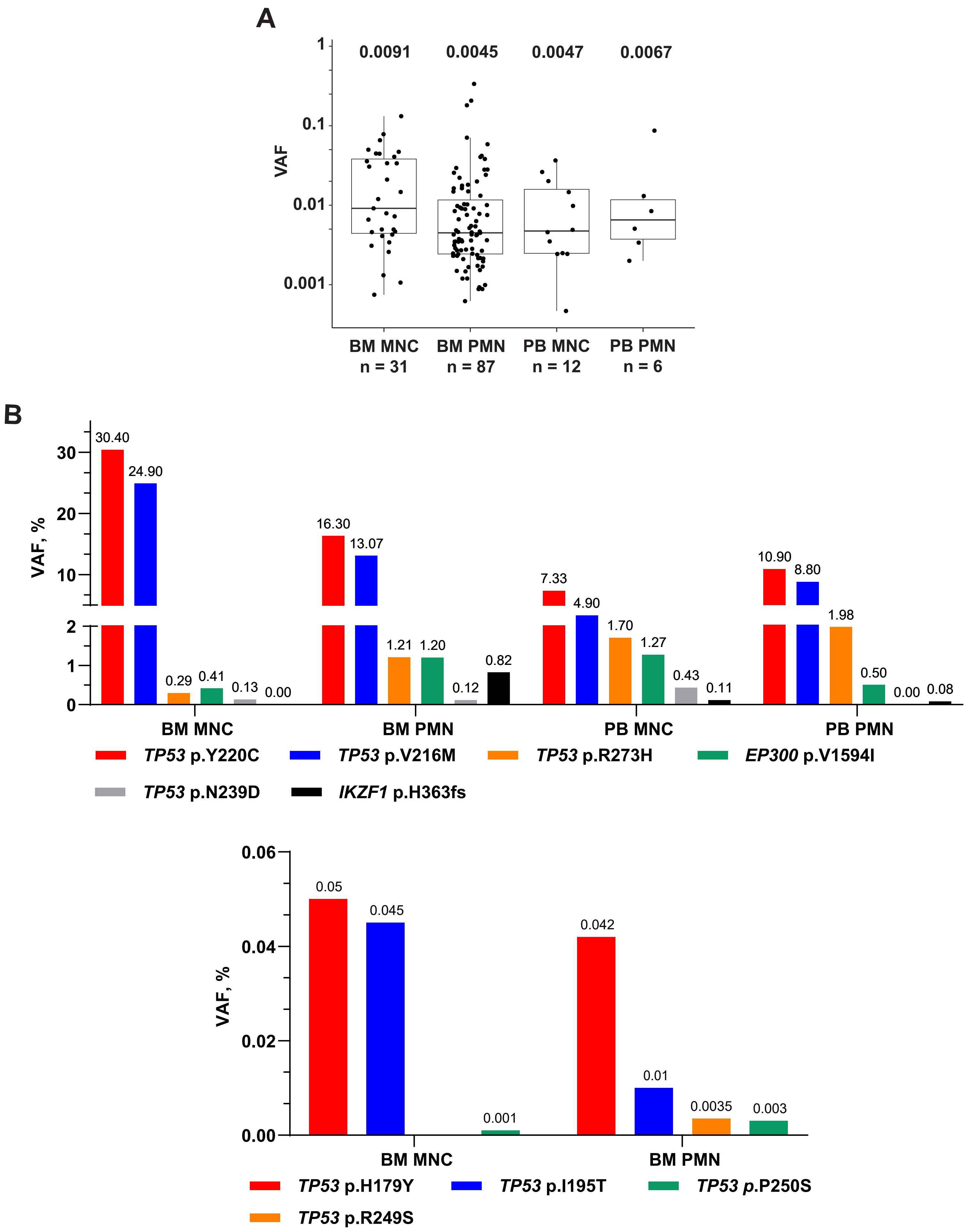


Supplementary figure 4

**A**

VAF

0 — 1

| Gene | Mutation | 1 | 2 | 3 | 4 | 5 | 6 | 7 | 8 |
|---|---|---|---|---|---|---|---|---|---|
| *ATM* | L459R | 0.486 | 0.567 | 0 | 0 | 0.377 | 0.42 | 0.395 | 0.28 |
| *KDM6A* | Intron | 0.462 | 0.422 | 0 | 0 | 0.355 | 0.178 | 0.385 | 0.178 |
| *TET2* | L1721W | 0.499 | 0.503 | 0 | 0 | 0.368 | 0.135 | 0.369 | 0.48 |
| *SETBP1* | V1101I | 0.492 | 0.495 | 0 | 0 | 0.331 | 0.121 | 0.381 | 0.234 |
| *ATM* | P1054R | 0.51 | 0.493 | 0 | 0 | 0.373 | 0.168 | 0.379 | 0.255 |
| *TP53* | P72R | 0.471 | 0.451 | 0.999 | 0.994 | 0.678 | 0.87 | 0.632 | 0.78 |
| *FLT3* | T227M | 0.994 | 0.996 | 0.499 | 0.512 | 0.86 | 0.649 | 0.868 | 1 |
| *CSF3R* | K704fs | 0.099 | 0 | 0 | 0 | 0 | 0 | 0 | 0 |
| *RUNX1* | R139fs | 0 | 0.736 | 0 | 0 | 0.5 | 0.21 | 0.51 | 0.314 |
| *CSF3R* | Q754* | 0 | 0.476 | 0 | 0 | 0.338 | 0.104 | 0.372 | 0.201 |
| *ASXL1* | S955fs | 0 | 0.479 | 0 | 0 | 0.329 | 0.092 | 0.33 | 0.194 |
| *SETD2* | P1962L | 0 | 0 | 0.995 | 0.971 | 0.312 | 0.772 | 0.275 | 0 |
| *BCL9* | 3'UTR | 0 | 0 | 0.429 | 0.506 | 0.158 | 0.345 | 0.147 | 0 |
| *KSR2* | Intron | 0 | 0 | 0.501 | 0.561 | 0.215 | 0.351 | 0.128 | 0 |
| *TNC* | L1594I | 0 | 0 | 0.481 | 0.488 | 0.204 | 0.324 | 0.125 | 0 |
| *SMARCA2* | R819W | 0 | 0 | 0.473 | 0.44 | 0.145 | 0.339 | 0.105 | 0 |
| *PRDM16* | P633L | 0 | 0 | 0.996 | 0.989 | 0.315 | 0.697 | 0.24 | 0 |
| *DYSF* | R1349L | 0 | 0 | 0.465 | 0.486 | 0.158 | 0.331 | 0.105 | 0 |
| *TET2* | I1762V | 0 | 0 | 0.497 | 0.52 | 0.152 | 0.37 | 0.123 | 0.294 |
| *TLX3* | 3'UTR | 0 | 0 | 0 | 0 | 0.02 | 0.023 | 0.033 | 0 |
| *KDM6A* | E36E | 0 | 0 | 0 | 0 | 0.023 | 0.023 | 0.029 | 0.0068 |
| *PAX5* | D127A | 0 | 0 | 0 | 0 | 0 | 0.011 | 0 | 0 |
| *KRAS* | G60V | 0 | 0 | 0 | 0 | 0 | 0 | 0.047 | 0.0051 |
| *KRAS* | G12V | 0 | 0 | 0 | 0 | 0 | 0 | 0.035 | 0.108 |
| *SALL4* | Intron | 0 | 0 | 0 | 0 | 0 | 0 | 0 | 0.157 |
| *BCORL1* | Intron | 0 | 0 | 0 | 0 | 0 | 0 | 0 | 0.25 |
| *SETD1B* | p.R1025Q | 0 | 0 | 0 | 0 | 0 | 0 | 0 | 0.265 |
| *TET2* | M1701I | 0 | 0 | 0 | 0 | 0 | 0 | 0 | 0.301 |
| *CBL* | c.e8-1 | 0 | 0 | 0 | 0 | 0 | 0 | 0 | 0.054 |
| *SETBP1* | P1130T | 0 | 0 | 0 | 0 | 0 | 0 | 0 | 0.249 |
| **Months from leukemia** | | -77.1 | 0 | 8.5 | 11.3 | 16.9 | 20.8 | 24.1 | 28.8 |

Patient germline | *CSF3R*-fs | Initiating | Donor specific | Acquired at the relapse

**B**

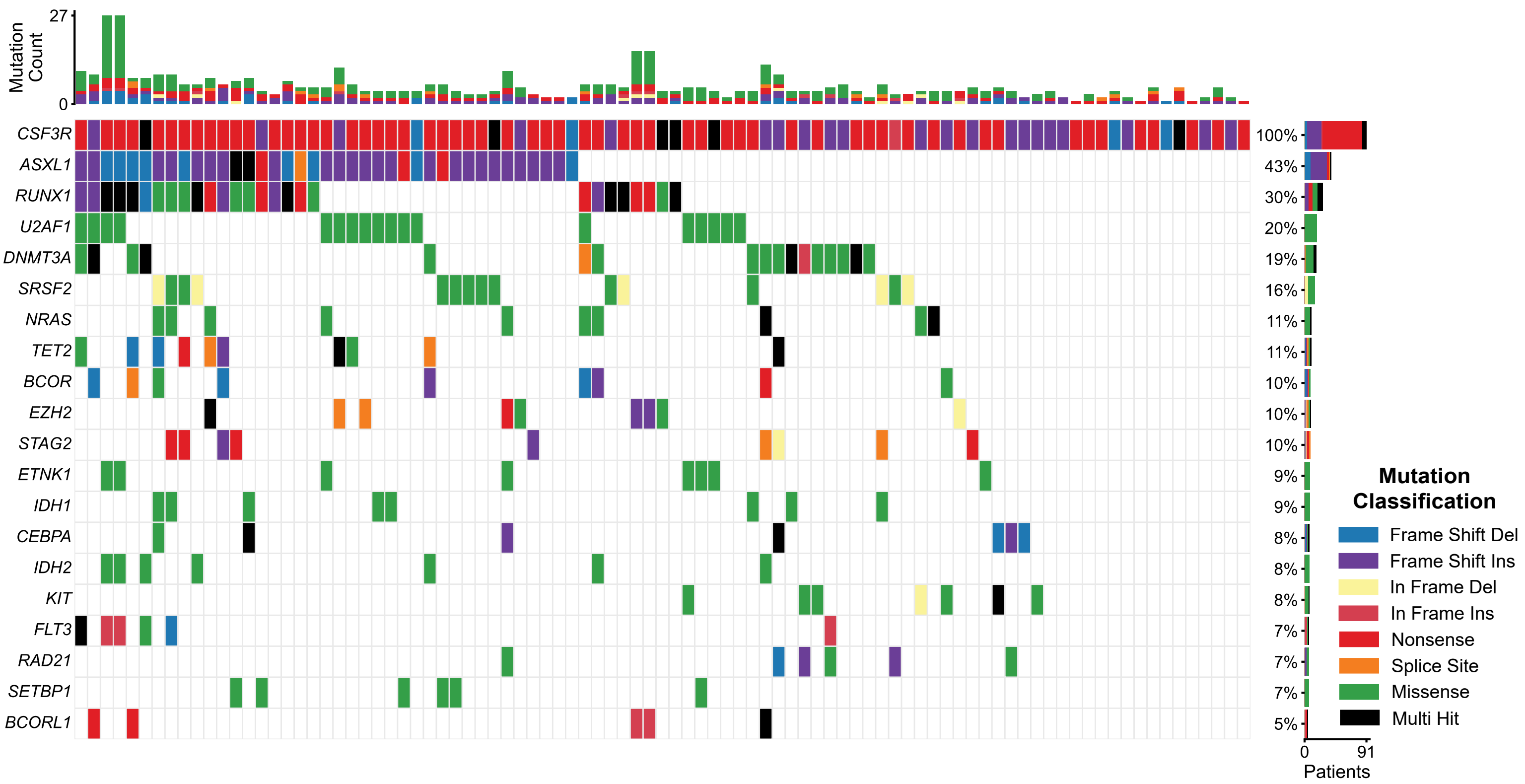


**Supplementary figure 5**

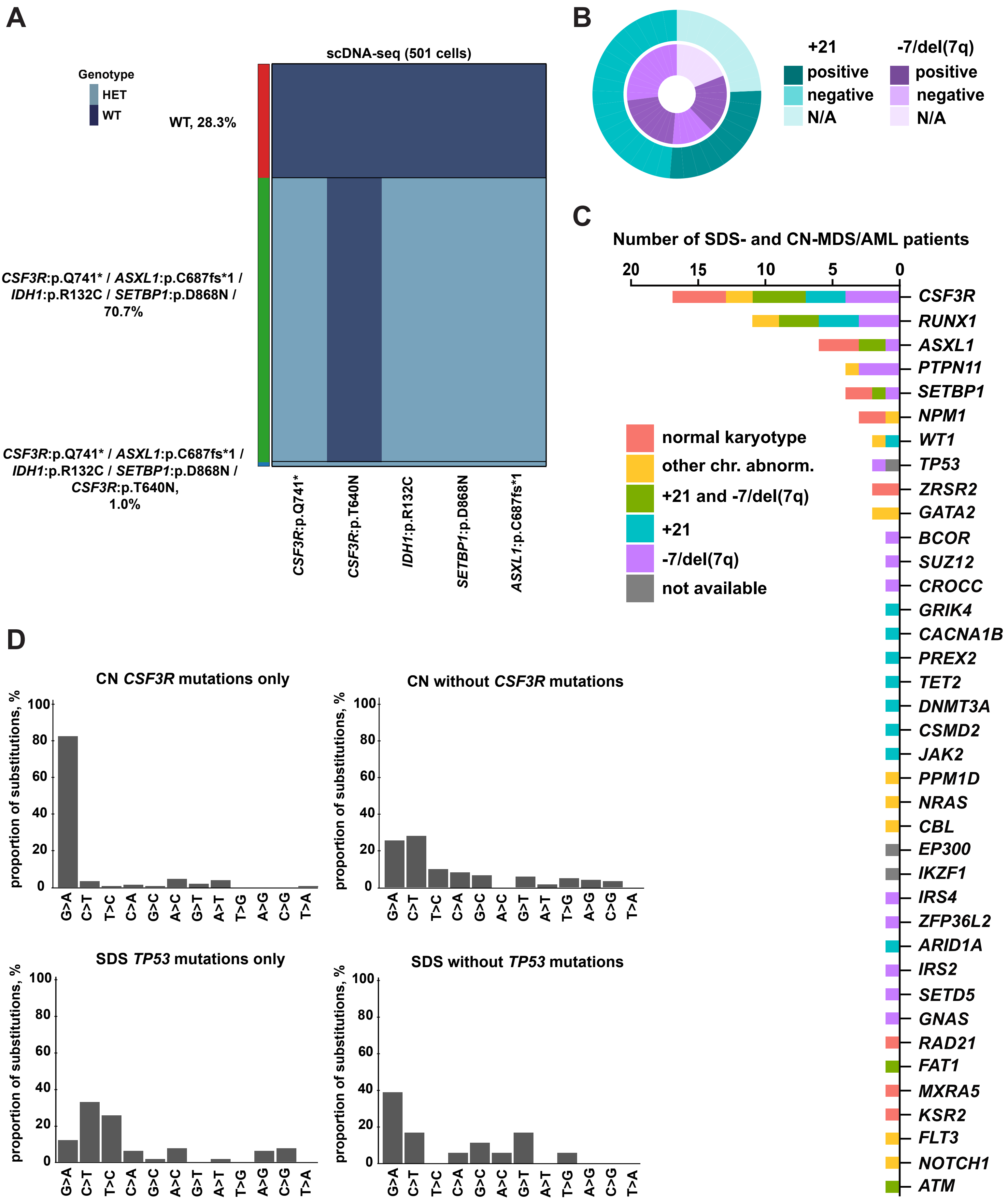


Supplementary figure 6
