## Supplementary_tables for "Deciphering the Dynamics of Leukemogenic Transformation from Clonal Hematopoiesis in Congenital Neutropenia"

**Supplementary table 1. Composition of the study cohort.**

| Neutropenia mutation | Neutropenia | Patients with samples before MDS/leukemia | MDS/ leukemia | After BMT (CN) | After BMT | Relapse |
|---|---|---|---|---|---|---|
| *ELANE* | 104 | 13 | 13 | 2 | 2 | 4 |
| *SBDS* | 54 | 6 | 4 | | | |
| neg_exome | 35 | | | | | |
| *HAX1* | 31 | 5 | 6 | | 1 | |
| *SRP54* | 14 | | | | | |
| *G6PC3* | 10 | | | | | |
| *JAGN1* | 6 | 1 | 1 | 1 | | |
| *CLPB* | 5 | | | | | |
| *GFI1* | 3 | | | | | |
| *TAFAZZIN* | 3 | | | | | |
| *TCIRG1* | 3 | | | | | |
| *VPS13B* | 3 | | | | | |
| *CXCR4* | 2 | | | | | |
| *SLC37A4* | 2 | | 1 | | | |
| *ADA2* | 1 | | | | | |
| *AP1B1* | 1 | | | | | |
| *CSF3R* | 1 | | | | | |
| *CXCR2* | 1 | | | | | |
| *GINS1* | 1 | | | | | |
| *SEC61A1* | 1 | | | | | |
| *WAS* | | 3 | 2 | | | |
| Total patients per category | **281** | **28** | **27** | **3** | **3** | **4** |
| Healthy: | | | | | | |
| Cord blood | 20 | | | | | |
| Healthy controls | 20 | | | | | |

**Supplementary table 2. Clinical, demographic, and genetic characteristics of the study cohort.**

| Patient_ID | Patient_sex | Sample_categories | Sample_count | Age at the last collection time, years | Observation period, years | Gene | Zygocity | Mutation |
|---|---|---|---|---|---|---|---|---|
| 1001 | f | progressive,leukemia,after_BMT,after_BMT,relapse,relapse,relapse,relapse | 8 | 40.73 | 8.70 | *ELANE* | heterozygous | NP_001963.1:p.Val101Met |
| 1002 | f | progressive,progressive,leukemia,relapse,relapse | 5 | 20.67 | 8.30 | *ELANE* | heterozygous | NP_001963.1:p.Ala233Pro, NP_001963.1:p.Val235Trpfs*1 |
| 1003 | m | leukemia | 1 | 18.35 | 0.00 | *ELANE* | heterozygous | NP_001963.1:Asn113Lys |
| 1004 | m | progressive,leukemia | 2 | 6.89 | 1.65 | *HAX1* | homozygous | NP_006109.2:p.Trp44* |
| 1005 | m | progressive,leukemia | 2 | 23.20 | 2.94 | *ELANE* | heterozygous | NP_001963.1:p.Ser126Leu |
| 1006 | m | progressive,leukemia,leukemia | 3 | 53.06 | 24.59 | *ELANE* | heterozygous | NP_001963.1:p.Val190_Phe199del |
| 1007 | f | progressive,leukemia | 2 | 21.17 | 12.35 | *ELANE* | heterozygous | NP_001963.1:p.Asp230fs |
| 1008 | f | progressive,leukemia | 2 | 11.32 | 10.64 | *HAX1* | homozygous | NP_006109.2:p.Trp44* |
| 1009 | m | progressive,progressive,progressive,leukemia | 4 | 15.98 | 5.52 | *HAX1* | homozygous | NP_006109.2:p.Trp44* |
| 1010 | f | progressive,leukemia | 2 | 16.78 | 8.37 | *ELANE* | heterozygous | NP_001963.1:p.Leu152Pro |
| 1011 | f | leukemia | 1 | 4.56 | 0.00 | *HAX1* | homozygous | NP_006109.2:p.Trp44* |
| 1012 | f | progressive,leukemia | 2 | 7.16 | 6.89 | *WAS* | hemizygous | NP_000368.1:p.Ser478Ile |
| 1013 | f | progressive,leukemia | 2 | 19.15 | 2.20 | *JAGN1* | homozygous | NM_032492.4:c.35_43del, NP_115881.3:p.Thr12_Gly14del |
| 1014 | m | progressive,leukemia | 2 | 2.93 | 2.80 | *ELANE* | heterozygous | NP_001963.1:p.Ala57Val |
| 1015 | m | progressive | 1 | 0.98 | 0.00 | *ELANE* | heterozygous | NP_001963.1:p.Gly214Arg |
| 1016 | m | MDS,MDS | 2 | 14.64 | 0.20 | *SBDS* | compound heterozygous | NM_016038.4:c.183_184delinsCT, NM_016038.4:258+2T>C |
| 1017 | f | neutropenia | 1 | 15.29 | 0.00 | *ELANE* | heterozygous | NP_001963.1:p.Cys223Alafs*17 |
| 1018 | m | neutropenia,neutropenia,neutropenia | 3 | 38.68 | 6.55 | *HAX1* | no_info | no_info |
| 1019 | m | neutropenia,neutropenia,neutropenia,neutropenia,neutropenia,neutropenia | 6 | 17.79 | 3.03 | *CLPB* | heterozygous | NM_030813.6:c.409G>T, NP_110440.1:p.Ala137Ser |
| 1020 | m | neutropenia,neutropenia,neutropenia,neutropenia,neutropenia,neutropenia,neutropenia,neutropenia,neutropenia,after_BMT_cn,after_BMT_cn,after_BMT_cn | 12 | 5.54 | 5.40 | *ELANE* | heterozygous | NP_001963.1:p.Val101Met |
| 1021 | m | neutropenia | 1 | 0.96 | 0.00 | *neg_exome* | | - |
| 1022 | f | neutropenia | 1 | 6.64 | 0.00 | *neg_exome* | | - |
| 1023 | f | neutropenia,neutropenia,neutropenia | 3 | 17.52 | 2.43 | *JAGN1* | homozygous | NP_115881.3:p.His44Tyr |
| 1024 | m | neutropenia,neutropenia,neutropenia,neutropenia,neutropenia | 5 | 22.41 | 4.25 | *ELANE* | heterozygous | NM_001972:c.337C>T;NP_001963.1:p.Ser126Leu |
| 1025 | f | neutropenia,neutropenia,neutropenia,neutropenia | 4 | 14.40 | 4.41 | *ELANE* | heterozygous | NP_001963.1:p.Ser126Leu |
| 1026 | f | neutropenia,neutropenia,neutropenia | 3 | 12.51 | 2.93 | *SBDS* | no_info | NM_016038.4:c.183_184delinsCT, NM_016038.4:258+2T>C |
| 1027 | m | neutropenia | 1 | 41.72 | 0.00 | *ELANE* | heterozygous | NM_001972.2:c.598-3C>G |

| | | | | | | | | |
|---|---|---|---|---|---|---|---|---|
| 1028 | f | neutropenia,neutropenia,neutropenia,neutropenia | 4 | 11.75 | 3.81 | *ELANE* | heterozygous | NM_006118.3:c.729G>A, NP_001963.1:p.Trp241* |
| 1029 | f | neutropenia,neutropenia,neutropenia,neutropenia,neutropenia | 5 | 7.98 | 4.24 | *ELANE* | heterozygous | NM_001972.4:c.136_141delTCCCTG, NP_001963.1:p.Ser46_Leu47del |
| 1030 | f | utropenia,neutropenia,neutropenia,neutropenia | 7 | 25.15 | 4.05 | *ELANE* | heterozygous | NM_001972.4:c.580+1G>A, NP_001963.1:p.V190_F199del |
| 1031 | f | neutropenia | 1 | 23.22 | 0.00 | *CSF3R* | compound heterozygous | NP_000751.1:p.Gln114*, NP_000751.1:p.Arg311Cys |
| 1032 | m | utropenia | 4 | 18.03 | 3.99 | *HAX1* | homozygous | NP_006109.2:p.Trp44* |
| 1033 | m | neutropenia,neutropenia | 2 | 3.99 | 0.07 | *ELANE* | heterozygous | NP_001963.1:p.Cys151Tyr |
| 1034 | f | neutropenia,neutropenia,neutropenia,neutropenia | 4 | 36.58 | 4.24 | *neg_exome* | | - |
| 1035 | f | neutropenia,neutropenia,neutropenia | 3 | 12.91 | 3.37 | *SBDS* | compound heterozygous | NM_016038.4:c.183_184delinsCT, NM_016038.4:258+2T>C |
| 1036 | f | neutropenia,neutropenia,neutropenia,neutropenia,neutropenia | 5 | 15.90 | 3.77 | *HAX1* | homozygous | NP_006109.2:p.Trp44* |
| 1037 | m | neutropenia,neutropenia,neutropenia,neutropenia | 4 | 14.44 | 3.31 | *TAFAZZIN* | no_info | no_info |
| 1038 | m | neutropenia,neutropenia,neutropenia | 3 | 8.75 | 3.68 | *HAX1* | homozygous | NM_006118.3:c.53+1G>A |
| 1039 | m | neutropenia,neutropenia,neutropenia,neutropenia,neutropenia | 5 | 11.04 | 4.29 | *ELANE* | heterozygous | NP_001963.1:p.Gly214Val |
| 1040 | f | neutropenia,neutropenia,neutropenia | 3 | 31.90 | 3.24 | *HAX1* | homozygous | NP_006109.2:p.Trp44* |
| 1041 | m | neutropenia,neutropenia,neutropenia,after_BMT_cn,after_BMT_cn,after_BMT_cn | 6 | 5.25 | 2.91 | *JAGN1* | homozygous | NP_115881.3:p.Glu21Asp |
| 1042 | m | neutropenia,neutropenia,neutropenia | 3 | 10.55 | 2.80 | *SBDS* | compound heterozygous | NM_016038.4:c.183_184delinsCT, NM_016038.4:258+2T>C |
| 1043 | f | neutropenia,neutropenia | 2 | 18.80 | 1.32 | *HAX1* | homozygous | NP_006109.2:p.Trp44* |
| 1044 | f | neutropenia,neutropenia | 2 | 17.97 | 0.98 | *G6PC3* | homozygous | NP_612396:p.Arg189* |
| 1045 | m | neutropenia,neutropenia | 2 | 18.28 | 1.13 | *ELANE* | heterozygous | NP_001963.1:p.Asp201Thrfs*1 |
| 1046 | f | neutropenia,neutropenia,neutropenia,neutropenia,neutropenia,neutropenia,neutropenia | 7 | 6.96 | 4.16 | *ELANE* | heterozygous | NP_001963.1:p.Val101Glu |
| 1047 | m | neutropenia | 1 | 17.60 | 0.00 | *ELANE* | heterozygous | NP_001963.1:p.Arg103Pro |
| 1048 | m | neutropenia,neutropenia | 2 | 11.09 | 2.53 | *ELANE* | heterozygous | NM_001972.4:c.580+5G>A, NP_001963.1:p.Val190_Phe199del |
| 1049 | m | neutropenia,neutropenia,neutropenia,neutropenia | 4 | 13.92 | 3.21 | *ELANE* | heterozygous | NP_001963.1:p.Gly221fs |
| 1050 | m | neutropenia,neutropenia,neutropenia,neutropenia | 4 | 17.85 | 3.45 | *AP1B1* | no_info | no_info |
| 1051 | f | neutropenia,neutropenia,neutropenia,neutropenia,neutropenia | 5 | 11.23 | 4.20 | *HAX1* | homozygous | NP_006109.2:p.Trp44* |
| 1052 | f | neutropenia | 1 | 29.43 | 0.00 | *neg_exome* | | - |

| | | | | | | | | |
|---|---|---|---|---|---|---|---|---|
| 1053 | m | neutropenia | 1 | 5.85 | 0.00 | *neg_exome* | | - |
| 1054 | m | after_BMT,after_BMT,after_BMT,after_BMT | 4 | 25.91 | 11.59 | *HAX1* | homozygous | NP_006109.2:p.Trp44* |
| 1055 | f | neutropenia,neutropenia,neutropenia,neutropenia,neutropenia | 5 | 15.34 | 4.20 | *HAX1* | homozygous | NP_006109.2:p.Trp44* |
| 1056 | f | neutropenia,neutropenia,neutropenia | 3 | 4.18 | 1.08 | *SBDS* | compound heterozygous | NM_016038.4:c.183_184delinsCT, NM_016038.4:258+2T>C |
| 1057 | m | neutropenia,neutropenia,neutropenia | 3 | 6.60 | 3.15 | *ELANE* | heterozygous | NP_001963.1:p.Phe199Leufs*16 |
| 1058 | m | neutropenia,neutropenia,neutropenia,neutropenia,neutropenia | 5 | 12.62 | 4.31 | *HAX1* | homozygous | NP_006109.2:p.Trp44* |
| 1059 | f | neutropenia,neutropenia,neutropenia,neutropenia | 4 | 17.83 | 3.90 | *ELANE* | heterozygous | NM_001972.4:c.580+1G>A |
| 1060 | f | neutropenia,neutropenia | 2 | 38.37 | 2.77 | *neg_exome* | | - |
| 1061 | m | neutropenia,neutropenia,neutropenia | 3 | 7.16 | 4.15 | *HAX1* | homozygous | NM_006118.3:c.504+112-_*413del, p.? |
| 1062 | f | neutropenia,neutropenia,neutropenia,neutropenia | 4 | 11.93 | 2.99 | *SBDS* | homozygous | NM_016038.4:258+2T>C |
| 1063 | f | neutropenia,neutropenia,neutropenia | 3 | 22.98 | 7.90 | *ELANE* | heterozygous | NM_001972:c.580+1G>A, NP_001963.1:p.Val190_Phe199del |
| 1064 | f | neutropenia,neutropenia | 2 | 34.02 | 5.46 | *ELANE* | heterozygous | NM_001972:c.580+1G>A, NP_001963.1:p.Val190_Phe199del |
| 1065 | m | neutropenia,neutropenia,neutropenia | 3 | 61.97 | 10.05 | *ELANE* | heterozygous | NM_001972:c.580+1G>A, NP_001963.1:p.Val190_Phe199del |
| 1066 | m | neutropenia | 1 | 6.04 | 0.00 | *HAX1* | homozygous | NP_006109.2:p.Trp44* |
| 1067 | m | neutropenia,neutropenia | 2 | 25.23 | 8.26 | *neg_exome* | | - |
| 1068 | f | neutropenia,neutropenia,neutropenia,neutropenia,neutropenia,neutropenia | 6 | 7.77 | 4.13 | *ELANE* | heterozygous | NP_001963.1:p.Val101Met |
| 1069 | f | neutropenia,neutropenia,neutropenia | 3 | 12.51 | 4.15 | *HAX1* | homozygous | NM_006118.3:c.504+112-_*413del,p.? |
| 1070 | m | neutropenia,neutropenia,neutropenia,neutropenia | 4 | 18.86 | 3.17 | *HAX1* | homozygous | NP_006109.2:p.Trp44* |
| 1071 | f | neutropenia | 1 | 32.75 | 0.00 | *neg_exome* | | - |
| 1072 | f | neutropenia,neutropenia | 2 | 6.02 | 1.82 | *ELANE* | heterozygous | NP_001963.1:p.Ser46Pro |
| 1073 | f | neutropenia | 1 | 15.78 | 0.00 | *neg_exome* | | - |
| 1074 | f | neutropenia,neutropenia,neutropenia,neutropenia,neutropenia,neutropenia,neutropenia,neutropenia,neutropenia,neutropenia,neutropenia,neutropenia | 12 | 16.73 | 4.35 | *ELANE* | heterozygous | NP_001963.1:p.Cys181Serfs*4 |
| 1075 | m | neutropenia,neutropenia | 2 | 15.72 | 1.63 | *SLC37A4* | homozygous | NM_000169.3:c.335G>A, NP_000160.1:p.Arg112His |
| 1076 | f | neutropenia | 1 | 15.23 | 0.00 | *ELANE* | heterozygous | NP_001963.1:p.Gly210Trp |
| 1077 | f | neutropenia,neutropenia,neutropenia | 3 | 9.90 | 1.38 | *SBDS* | no_info | no_info |
| 1078 | f | neutropenia | 1 | 15.31 | 0.00 | *ELANE* | heterozygous | NP_001963.1:p.Cys208Gly |
| 1079 | m | neutropenia,neutropenia | 2 | 12.41 | 2.06 | *neg_exome* | | - |

| | | | | | | | | |
|---|---|---|---|---|---|---|---|---|
| 1080 | f | neutropenia,neutropenia | 2 | 36.47 | 3.58 | *ELANE* | heterozygous | NP_001963.1:p.Ser126Leu |
| 1081 | m | neutropenia,neutropenia,neutropenia,neutropenia,neutropenia | 5 | 6.73 | 3.96 | *SBDS* | compound heterozygous | NM_016038.4:c.183_184delinsCT, NM_016038.4:258+2T>C |
| 1082 | m | neutropenia,neutropenia,neutropenia,neutropenia | 4 | 35.55 | 3.56 | *G6PC3* | homozygous | NM_138387.3:c.778G>C, NP_612396.1:p.Gly260Arg |
| 1083 | f | neutropenia | 1 | 1.46 | 0.00 | *neg_exome* | | - |
| 1084 | f | neutropenia,neutropenia,neutropenia,neutropenia,neutropenia | 5 | 14.75 | 4.27 | *ELANE* | heterozygous | NM_001972.2:c.597+5G, NP_001963.1:p.Val190_Phe199del |
| 1085 | m | neutropenia,neutropenia,neutropenia | 3 | 7.71 | 3.67 | *SBDS* | compound heterozygous | NM_016038.4:258+2T>C, het. loss of SBDS exon 2 |
| 1086 | m | neutropenia | 1 | 7.75 | 0.00 | *HAX1* | homozygous | NP_006109.2:p.Trp44* |
| 1087 | m | neutropenia,neutropenia | 2 | 19.20 | 1.04 | *JAGN1* | no_info | no_info |
| 1088 | f | neutropenia,neutropenia,neutropenia,neutropenia | 4 | 7.28 | 3.65 | *SLC37A4* | heterozygous, verification needed | NM_001164277.2:c.1042_1043del, NP_001157749.1:p.Leu348fs |
| 1089 | f | neutropenia,neutropenia | 2 | 22.91 | 2.83 | *G6PC3* | homozygous | NP_612396:p.Gly260Arg |
| 1090 | m | neutropenia,neutropenia,neutropenia | 3 | 6.55 | 3.51 | *ELANE* | no_info | no_info |
| 1091 | m | neutropenia | 1 | 5.67 | 0.00 | *SRP54* | heterozygous | NM_003136.4:c.349_351del, NP_003127.1:p.Thr117del |
| 1092 | f | neutropenia | 1 | 12.27 | 0.00 | *ELANE* | heterozygous | NM_001972.2:p.Gly203Cys |
| 1093 | m | progressive,progressive,leukemia | 3 | 24.44 | 1.52 | *SBDS* | compound heterozygous | NM_016038.4:c.183_184delinsCT, NM_016038.4:258+2T>C |
| 1094 | f | neutropenia,neutropenia,neutropenia,neutropenia | 4 | 25.67 | 3.41 | *HAX1* | homozygous | NP_006109.2:p.Trp44* |
| 1095 | m | neutropenia | 1 | 25.07 | 0.00 | *HAX1* | homozygous | NP_006109.2:p.Trp44Ter |
| 1096 | f | neutropenia,neutropenia | 2 | 1.23 | 0.38 | *ELANE* | heterozygous | NP_001963.1:p.Ala57Thr |
| 1097 | m | neutropenia | 1 | 12.12 | 0.00 | *ELANE* | no_info | no_info |
| 1098 | m | neutropenia,neutropenia,neutropenia,neutropenia | 4 | 13.95 | 3.72 | *G6PC3* | no_info | no_info |
| 1099 | f | neutropenia,neutropenia | 2 | 28.82 | 2.95 | *HAX1* | homozygous | NP_006109.2:p.Trp44* |
| 1100 | m | neutropenia,neutropenia,neutropenia,neutropenia,neutropenia | 5 | 13.46 | 3.30 | *SBDS* | compound heterozygous | NM_016038.4:c.183_184delinsCT, NM_016038.4:258+2T>C |
| 1101 | f | neutropenia | 1 | 19.87 | 0.00 | *SBDS* | compound heterozygous | NM_016038.4:c.183_184delinsCT, NM_016038.4:258+2T>C |
| 1102 | m | neutropenia,neutropenia,neutropenia,neutropenia | 4 | 20.99 | 3.23 | *ELANE* | heterozygous | NP_001963.1:p.Val190_Phe199del |
| 1103 | m | neutropenia,neutropenia,neutropenia,neutropenia | 4 | 20.18 | 2.85 | *G6PC3* | homozygous | NM_138387.4:c.758G>A, NP_612396.1:p.Arg253His |
| 1104 | m | neutropenia | 1 | 23.50 | 0.00 | *ELANE* | heterozygous | NM_001972.4:c.580+1 G>A, NP_001963.1:p.Val190_Phe199del |
| 1105 | m | neutropenia | 1 | 27.86 | 0.00 | *ELANE* | heterozygous | NM_001972.4:c.580+1 G>A, NP_001963.1:p.Val190_Phe199del |
| 1106 | m | neutropenia | 1 | 2.47 | 0.00 | *TCIRG1* | no_info | no_info |
| 1107 | m | neutropenia | 1 | 0.37 | 0.00 | *SRP54* | no_info | no_info |
| 1108 | m | neutropenia | 1 | 0.93 | 0.00 | *SRP54* | heterozygous | NM_003136.4:c.349_351del, NP_003127.1:p.Thr117del |
| 1109 | f | neutropenia | 1 | 14.43 | 0.00 | *CLPB* | no_info | no_info |

| | | | | | | | | |
|---|---|---|---|---|---|---|---|---|
| 1110 | f | neutropenia,neutropenia | 2 | 5.72 | 0.30 | *CLPB* | heterozygous | NM_030813.6:c.1222A>G, NP_110440.1:p.Arg408Gly |
| 1111 | f | neutropenia,neutropenia,neutropenia,neutropenia | 4 | 8.22 | 3.09 | *SRP54* | heterozygous | NP_115881.3:p.Thr117del |
| 1112 | f | neutropenia | 1 | 1.03 | 0.00 | *ELANE* | heterozygous | NM_001972:c.169G>A, NP_001963.1:p.Ala57Thr |
| 1113 | m | neutropenia,neutropenia,neutropenia | 3 | 16.35 | 1.40 | *ELANE* | heterozygous | NP_001963.1:Ala79Valfs*9 |
| 1114 | m | neutropenia,neutropenia | 2 | 18.05 | 0.53 | *ELANE* | heterozygous | NP_001963.1:p.Val197Alafs*2 |
| 1115 | m | neutropenia,neutropenia,neutropenia,neutropenia,neutropenia | 5 | 14.57 | 3.90 | *ELANE* | heterozygous | NM_001972.2:c.241_246delCGGGTG, NP_001963.1:p.Arg81_Val82del |
| 1116 | m | neutropenia,neutropenia,neutropenia,neutropenia | 4 | 6.32 | 4.56 | *SRP54* | heterozygous | NP_115881.3:p.Thr117del |
| 1117 | f | neutropenia,neutropenia,neutropenia,neutropenia | 4 | 8.22 | 3.09 | *SRP54* | heterozygous | NP_115881.3:p.Thr117del |
| 1118 | f | neutropenia,neutropenia | 2 | 17.21 | 1.19 | *SBDS* | compound heterozygous | NM_016038.4:c.183_184delinsCT, NM_016038.4:258+2T>C |
| 1119 | m | neutropenia,neutropenia,neutropenia,neutropenia | 4 | 7.36 | 3.97 | *SRP54* | heterozygous | NP_115881.3:p.Thr117del |
| 1120 | m | neutropenia,neutropenia,neutropenia | 3 | 17.76 | 2.05 | *SBDS* | compound heterozygous | NM_016038.4:c.183_184delinsCT, NM_016038.4:258+2T>C |
| 1121 | f | neutropenia,neutropenia,neutropenia,neutropenia,neutropenia | 5 | 4.55 | 3.10 | *SBDS* | compound heterozygous | NM_016038.4:c.183_184delinsCT, NM_016038.4:258+2T>C |
| 1122 | f | neutropenia,neutropenia | 2 | 12.60 | 1.46 | *ELANE* | heterozygous | NP_001963.1:p.Ser126Leu |
| 1123 | f | neutropenia | 1 | 65.12 | 0.00 | *ELANE* | heterozygous | NP_001963.1:p.Trp241Gly |
| 1124 | m | neutropenia | 1 | 32.44 | 0.00 | *ELANE* | heterozygous | NP_001963.1:p.Trp241Gly |
| 1125 | m | neutropenia,neutropenia,neutropenia | 3 | 18.10 | 1.30 | *neg_exome* | | - |
| 1126 | m | neutropenia,neutropenia,neutropenia | 3 | 24.35 | 1.66 | *neg_exome* | | - |
| 1127 | f | neutropenia | 1 | 16.23 | 0.00 | *SBDS* | no_info | no_info |
| 1128 | m | neutropenia,neutropenia,neutropenia,neutropenia,neutropenia | 5 | 5.98 | 3.61 | *ELANE* | heterozygous | NP_001963.1:p.Ile120Phe |
| 1129 | f | neutropenia | 1 | 36.42 | 0.00 | *G6PC3* | homozygous | NP_612396.1:p.Met116Lys |
| 1130 | f | neutropenia,neutropenia,neutropenia | 3 | 18.74 | 2.75 | *ELANE* | heterozygous | NP_001963.1:p.Ser126Leu |
| 1131 | f | neutropenia | 1 | 9.34 | 0.00 | *ELANE* | heterozygous | NP_001963.1:p.Trp156Cys |
| 1132 | m | neutropenia,neutropenia,neutropenia,neutropenia | 4 | 13.20 | 3.52 | *ELANE* | heterozygous | NP_001963.1:p.Gln194* |
| 1133 | f | neutropenia,neutropenia,neutropenia,neutropenia | 4 | 7.91 | 3.01 | *GINS1* | compound heterozygous | NM_021067.5:c.247C>T, NP_066545.3:p.Arg83Cys; NM_021067.5:c.29T>G, NP_066545.3:p.Ile10Ser |
| 1134 | f | neutropenia,neutropenia,neutropenia,neutropenia | 4 | 23.62 | 4.69 | *VPS13B* | no_info | no_info |
| 1135 | m | neutropenia,neutropenia,neutropenia,neutropenia | 4 | 20.46 | 4.69 | *VPS13B* | no_info | no_info |
| 1136 | f | neutropenia,neutropenia,neutropenia | 3 | 7.72 | 1.55 | *SBDS* | compound heterozygous | NM_016038.4:c.183_184delinsCT, NM_016038.4:258+2T>C |

| | | | | | | | | |
|---|---|---|---|---|---|---|---|---|
| 1137 | m | neutropenia,neutropenia,neutropenia,neutropenia | 4 | 5.57 | 2.82 | *SBDS* | compound heterozygous | NM_016038.4:c.183_184delinsCT, NM_016038.4:258+2T>C |
| 1138 | f | neutropenia,neutropenia,neutropenia,neutropenia | 4 | 3.58 | 3.33 | *ELANE* | heterozygous | NM_006118.3:c.597+5G>A |
| 1139 | f | neutropenia | 1 | 17.11 | 0.00 | *G6PC3* | homozygous | NM_138387.4:c.758G>A, NP_612396.1:p.Arg253His |
| 1140 | f | neutropenia,neutropenia | 2 | 17.80 | 0.98 | *neg_exome* | | - |
| 1141 | m | neutropenia | 1 | 34.33 | 0.00 | *ELANE* | heterozygous | NP_001963.1:p.Ser126Trp |
| 1142 | f | neutropenia | 1 | 0.26 | 0.00 | *ELANE* | heterozygous | NP_001963.1:p.Ser126Trp |
| 1143 | f | neutropenia | 1 | 12.78 | 0.00 | *HAX1* | homozygous | NP_006109.2:p.Val144Glyfs*5 |
| 1144 | m | neutropenia | 1 | 3.26 | 0.00 | *ELANE* | heterozygous | NM_006118.3:c.585delC, NP_001963.1:p.Gly196Alafs*16 |
| 1145 | m | neutropenia,neutropenia,neutropenia | 3 | 18.60 | 3.29 | *ELANE* | heterozygous | NP_001963.1:p.Pro110Arg |
| 1146 | f | neutropenia,neutropenia | 2 | 16.85 | 3.12 | *neg_exome* | | - |
| 1147 | f | neutropenia,neutropenia | 2 | 21.00 | 3.12 | *neg_exome* | | - |
| 1148 | f | progressive | 1 | 15.72 | 0.00 | *WAS* | hemizygous | NP_000368.1:p.Ile331Met |
| 1149 | m | neutropenia | 1 | 22.06 | 0.00 | *TCIRG1* | no_info | - |
| 1150 | f | progressive,progressive | 2 | 46.73 | 2.02 | *ELANE* | heterozygous | NP_001963.1:p.Ser126Leu |
| 1151 | m | neutropenia | 1 | 37.73 | 0.00 | *SRP54* | no_info | no_info |
| 1152 | f | neutropenia,neutropenia | 2 | 28.07 | 1.26 | *VPS13B* | no_info | no_info |
| 1153 | f | neutropenia,neutropenia | 2 | 2.40 | 1.51 | *ELANE* | heterozygous | NM_006118.3:c.544delC, NP_001963.1:p.Arg182Valfs*9 |
| 1154 | m | neutropenia,neutropenia,neutropenia,neutropenia,neutropenia | 5 | 9.80 | 3.06 | *HAX1* | homozygous | NP_006109.2:p.Trp44* |
| 1155 | m | neutropenia | 1 | 17.78 | 0.00 | *GFI1* | heterozygous | NM_001127215.3:c.1145A>G, NP_005254.2:p.Asn382Ser |
| 1156 | f | neutropenia | 1 | 15.82 | 0.00 | *ADA2* | compound heterozygous | NM_001282225.2:c.506G>A, NP_001269154.1:p.Arg169Gln; NM_001282225.2:c.973-2A>G |
| 1157 | m | neutropenia,neutropenia,neutropenia,neutropenia | 4 | 17.30 | 3.01 | *JAGN1* | homozygous | NM_032492.4:c.59G>A, NP_115881.3:p.Arg20Gln |
| 1158 | m | progressive | 1 | 12.39 | 0.00 | *ELANE* | no_info | no_info |
| 1159 | m | progressive,progressive,leukemia | 3 | 17.63 | 14.03 | *ELANE* | no_info | no_info |
| 1160 | m | progressive,MDS,leukemia | 3 | 7.84 | 4.97 | *ELANE* | heterozygous | NP_001963.1:p.Tyr228* |
| 1161 | m | progressive,progressive | 2 | 10.32 | 8.03 | *SBDS* | no_info | no_info |
| 1162 | f | progressive,MDS | 2 | 20.30 | 1.55 | *WAS* | no_info | no_info |
| 1163 | f | neutropenia | 1 | 10.38 | 0.00 | *TCIRG1* | no_info | no_info |
| 1164 | f | neutropenia | 1 | 38.84 | 0.00 | *neg_exome* | | - |
| 1165 | m | neutropenia | 1 | 17.45 | 0.00 | *ELANE* | heterozygous | NP_001963.1:p.Pro139Leu |
| 1166 | f | progressive,progressive,progressive | 3 | 21.34 | 1.28 | *SBDS* | no_info | no_info |
| 1167 | m | neutropenia,neutropenia,neutropenia | 3 | 35.49 | 6.79 | *JAGN1* | homozygous | NP_115881.3:p.Glu21Asp |
| 1168 | f | neutropenia | 1 | 7.44 | 0.00 | *ELANE* | heterozygous | NM_001972.2:c.163T>C, NP_001963.1:p.Cys55Arg |
| 1169 | f | neutropenia | 1 | 13.56 | 0.00 | *ELANE* | homozygous | NM_001972.2:c.659G>A, NP_001963.1:p.Arg220Gln |
| 1170 | m | neutropenia,neutropenia,neutropenia | 3 | 12.08 | 2.46 | *HAX1* | homozygous | NP_006109.2:p.Trp44* |
| 1171 | m | neutropenia | 1 | 15.89 | 0.00 | *ELANE* | homozygous | NM_001972.2:c.659G>A, NP_001963.1:p.Arg220Gln |

| | | | | | | | | |
|---|---|---|---|---|---|---|---|---|
| 1172 | f | neutropenia | 1 | 13.90 | 0.00 | *ELANE* | homozygous | NM_001972.2:c.659G>A, NP_001963.1:p.Arg220Gln |
| 1173 | m | neutropenia,neutropenia,neutropenia | 3 | 7.85 | 2.15 | *ELANE* | heterozygous | NM_001972.2:c.659G>A, NP_001963.1:p.Arg220Gln |
| 1174 | m | neutropenia,neutropenia,neutropenia | 3 | 9.76 | 2.91 | *ELANE* | heterozygous | NP_001963.1:p.Ile120Phe |
| 1175 | m | neutropenia | 1 | 2.86 | 0.00 | *ELANE* | heterozygous | NM_001972.2:c.597+5G>A |
| 1176 | m | neutropenia,neutropenia | 2 | 12.26 | 2.11 | *SBDS* | no_info | no_info |
| 1177 | f | neutropenia | 1 | 17.07 | 0.00 | *neg_exome* | | - |
| 1178 | m | after_BMT,relapse | 2 | 29.00 | 1.86 | *ELANE* | heterozygous | NP_001963.1:Ser67Trp |
| 1179 | f | neutropenia | 1 | 9.34 | 0.00 | *neg_exome* | | - |
| 1180 | f | neutropenia,neutropenia,neutropenia | 3 | 31.96 | 2.17 | *ELANE* | heterozygous | NM_001972.4:c.659G>A, NP_001963.1:p.Arg220Gln |
| 1181 | f | neutropenia,neutropenia | 2 | 13.92 | 2.03 | *SBDS* | compound heterozygous | NM_016038.4:c.183_184delinsCT, NM_016038.4:258+2T>C |
| 1182 | m | neutropenia | 1 | 3.66 | 0.00 | *HAX1* | homozygous | NP_006109.2:p.Trp44* |
| 1183 | f | neutropenia | 1 | 12.43 | 0.00 | *ELANE* | heterozygous | NP_001963.1:p.Ser126Leu |
| 1184 | m | neutropenia,neutropenia | 2 | 36.46 | 5.97 | *ELANE* | heterozygous | NP_001963.1:p.Ile120Phe |
| 1185 | m | neutropenia | 1 | 17.92 | 0.00 | *ELANE* | heterozygous | NP_001963.1:p.Val190_Phe199del |
| 1186 | m | neutropenia | 1 | 16.90 | 0.00 | *ELANE* | heterozygous | NM_001972.2: c.140T>G, NP_001963.1:p.Leu47Arg |
| 1187 | m | neutropenia | 1 | 37.81 | 0.00 | *ELANE* | heterozygous | NP_001963.1:p.Ala57Thr |
| 1188 | m | neutropenia | 1 | 27.98 | 0.00 | *ELANE* | heterozygous | NP_001963.1:p.Phe199Leufs*8 |
| 1189 | m | neutropenia | 1 | | 0.00 | *ELANE* | heterozygous | NP_001963.1:p.Ser123Leu |
| 1190 | f | neutropenia | 1 | 6.30 | 0.00 | *ELANE* | no_info | no_info |
| 1191 | m | progressive,leukemia | 2 | 11.10 | 1.86 | *HAX1* | homozygous | NP_006109.2:p.Trp44* |
| 1192 | f | neutropenia | 1 | 7.03 | 0.00 | *SBDS* | no_info | no_info |
| 1193 | m | neutropenia | 1 | 8.85 | 0.00 | *TAFAZZIN* | no_info | no_info |
| 1194 | f | neutropenia,neutropenia,neutropenia | 3 | 36.93 | 1.78 | *ELANE* | heterozygous | NM_001972.2:c.455T>C, NP_001963.1:p.Leu152Pro |
| 1195 | m | neutropenia,neutropenia | 2 | 31.99 | 2.77 | *CLPB* | heterozygous | NM_030813.6:c.569G>T, NP_110440.1:p.Gly190Val |
| 1196 | f | neutropenia,neutropenia | 2 | 1.97 | 1.32 | *neg_exome* | | - |
| 1197 | m | neutropenia,neutropenia | 2 | 8.23 | 1.99 | *SBDS* | compound heterozygous | NM_016038.4:c.183_184delinsCT, NM_016038.4:258+2T>C |
| 1198 | m | neutropenia | 1 | 6.68 | 0.00 | *GFI1* | no_info | no_info |
| 1199 | f | leukemia | 1 | 11.37 | 0.00 | *SLC37A4* | compound heterozygous | NM_001164277.1:c.1042_1043delCT, NP_001157749.1:p.Leu348fs, NM_001164277.2:c.985+1G>A |
| 1200 | m | progressive,leukemia | 2 | 10.63 | 0.34 | *ELANE* | heterozygous | NP_001963.1:p.Gly214Arg |
| 1201 | m | neutropenia | 1 | 2.55 | 0.00 | *ELANE* | heterozygous | NM_001972.2:c.457G>C, NP_001963.1:p.Ala153Pro |
| 1202 | m | progressive | 1 | 44.93 | 0.00 | *SBDS* | compound heterozygous | NM_016038.4:c.183_184delinsCT, NM_016038.4:258+2T>C |
| 1203 | f | neutropenia,neutropenia,neutropenia | 3 | 8.21 | 2.03 | *SBDS* | compound heterozygous | NM_016038.4:c.183_184delinsCT, NM_016038.4:258+2T>C |
| 1204 | m | neutropenia,neutropenia,neutropenia | 3 | 8.91 | 1.85 | *SBDS* | compound heterozygous | NM_016038.4:c.183_184delinsCT, NM_016038.4:258+2T>C |
| 1205 | m | neutropenia,neutropenia,neutropenia | 3 | 10.32 | 2.20 | *GFI1* | heterozygous | NM_005263.3:c.1145A>G, NP_005254.2:p.Asn382Ser |
| 1206 | m | neutropenia | 1 | 4.95 | 0.00 | *ELANE* | heterozygous | NM_001972:c.597+5G>A, NP_001963.1:p.Val190_Phe199del |

| | | | | | | | | |
|---|---|---|---|---|---|---|---|---|
| 1207 | f | neutropenia | 1 | 9.06 | 0.00 | *neg_exome* | | - |
| 1208 | m | neutropenia,neutropenia | 2 | 13.72 | 2.02 | *neg_exome* | | - |
| 1209 | f | neutropenia | 1 | 13.28 | 0.00 | *neg_exome* | | - |
| 1210 | f | neutropenia,neutropenia,neutropenia,neutropenia | 4 | 4.89 | 3.61 | *SEC61A1* | homozygous | NM_013336.4:c.275A>G, NP_037468.1:p.Gln92Arg |
| 1211 | m | progressive,progressive,MDS | 3 | 11.09 | 1.34 | *SBDS* | compound heterozygous | NM_016038.4:c.183_184delinsCT, NM_016038.4:258+2T>C |
| 1212 | f | neutropenia | 1 | 10.09 | 0.00 | *ELANE* | heterozygous | NP_001963.1:p.Pro139Leu |
| 1213 | m | leukemia | 1 | 55.04 | 0.00 | *ELANE* | heterozygous | NP_001963.1:p.Val190_Phe199del |
| 1214 | m | neutropenia,neutropenia,neutropenia | 3 | 9.22 | 1.57 | *SBDS* | compound heterozygous | NM_016038.4:c.356G>A, NP_057122.2:p.Cys119Tyr; NM_016038.4:258+2T>C |
| 1215 | f | neutropenia,neutropenia,neutropenia | 3 | 2.61 | 2.12 | *HAX1* | homozygous | NM_006118.4:c.505-1G>C |
| 1216 | m | neutropenia | 1 | 49.29 | 0.00 | *neg_exome* | | - |
| 1217 | m | neutropenia | 1 | 42.76 | 0.00 | *neg_exome* | | - |
| 1218 | f | neutropenia | 1 | 20.53 | 0.00 | *neg_exome* | | - |
| 1219 | f | neutropenia | 1 | 26.66 | 0.00 | *neg_exome* | | - |
| 1220 | f | neutropenia | 1 | 27.82 | 0.00 | *CXCR2* | | no_info |
| 1221 | m | neutropenia | 1 | 12.99 | 0.00 | *ELANE* | heterozygous | NM_001972.2:c.597+1G>A, NP_001963.1:p.Val190_Phe199del |
| 1222 | m | neutropenia | 1 | 14.20 | 0.00 | *HAX1* | homozygous | NP_006109.2:p.Trp44* |
| 1223 | m | neutropenia,neutropenia | 2 | 15.16 | 1.76 | *SBDS* | no_info | no_info |
| 1224 | f | relapse | 1 | 26.28 | 0.00 | *ELANE* | heterozygous | NM_001972.2:c.377C>T, NP_001963.1:p.Ser126Leu |
| 1225 | f | neutropenia | 1 | 16.83 | 0.00 | *ELANE* | no_info | no_info |
| 1226 | m | progressive,leukemia,leukemia | 3 | 15.00 | 6.80 | *HAX1* | homozygous | NP_006109.2:p.Trp44* |
| 1227 | m | neutropenia,neutropenia | 2 | 4.09 | 1.89 | *SBDS* | compound heterozygous | NM_016038.4:258+2T>C, NM_016038.4:c.127G>T, NP_057122.2:p.Val43Leu |
| 1228 | m | neutropenia | 1 | 31.57 | 0.00 | *neg_exome* | | - |
| 1229 | m | neutropenia | 1 | 55.92 | 0.00 | *ELANE* | heterozygous | NP_001963.1:p.Val190_Phe199del |
| 1231 | m | neutropenia | 1 | 49.68 | 0.00 | *ELANE* | heterozygous | NP_001963.1:p.Ser126Leu |
| 1232 | f | neutropenia | 1 | 47.22 | 0.00 | *ELANE* | heterozygous | NP_001963.1:p.Val190_Phe199del |
| 1233 | f | neutropenia | 1 | 50.93 | 0.00 | *ELANE* | heterozygous | NP_001963.1:p.Gly196Profs*17 |
| 1234 | f | neutropenia,neutropenia | 2 | 28.33 | 0.32 | *CXCR4* | heterozygous | NM_003467.3:c.1000C>T, NP_003458.1:p.Arg334* |
| 1235 | f | neutropenia,neutropenia,neutropenia | 3 | 47.80 | 2.03 | *ELANE* | heterozygous | NP_001963.1:p.Val239* |
| 1236 | m | neutropenia | 1 | 12.64 | 0.00 | *neg_exome* | | - |
| 1237 | m | neutropenia | 1 | 3.70 | 0.00 | *HAX1* | homozygous | NP_006109.2:p.Trp44* |
| 1238 | f | neutropenia,neutropenia,neutropenia | 3 | 9.32 | 2.00 | *SBDS* | no_info | no_info |

| | | | | | | | | |
|---|---|---|---|---|---|---|---|---|
| 1239 | m | neutropenia,neutropenia,neutropenia | 3 | 3.85 | 1.99 | *ELANE* | heterozygous | NP_001963.1:p.Ile120Phe |
| 1240 | m | neutropenia,neutropenia,neutropenia | 3 | 7.90 | 2.11 | *SBDS* | compound heterozygous | NM_016038.4:c.183_184delinsCT, NM_016038.4:258+2T>C |
| 1241 | m | neutropenia,neutropenia,neutropenia | 3 | 10.31 | 2.11 | *SBDS* | compound heterozygous | NM_016038.4:c.183_184delinsCT, NM_016038.4:258+2T>C |
| 1242 | f | neutropenia,neutropenia,neutropenia | 3 | 3.41 | 1.99 | *SBDS* | no_info | no_info |
| 1243 | f | neutropenia,neutropenia | 2 | 5.79 | 1.02 | *SRP54* | heterozygous | NM_003136.4:c.349_351del, NP_003127.1:p.Thr117del |
| 1244 | m | neutropenia | 1 | 6.42 | 0.00 | *HAX1* | homozygous | NP_006109.2:p.Trp44* |
| 1245 | f | neutropenia,neutropenia,neutropenia | 3 | 12.97 | 2.03 | *ELANE* | no_info | no_info |
| 1246 | f | neutropenia,neutropenia,neutropenia | 3 | 1.71 | 1.66 | *ELANE* | heterozygous | NP_001963.1:p.Cys223* |
| 1247 | m | neutropenia | 1 | 4.38 | 0.00 | *SBDS* | compound heterozygous | NM_016038.4:c.624+1G>A, NM_016038.4:258+2T>C |
| 1248 | m | neutropenia,neutropenia | 2 | 14.52 | 1.19 | *ELANE* | heterozygous | NM_001972.2:c.377C>T, NP_001963.1:p.Ser126Leu |
| 1249 | m | neutropenia,neutropenia | 2 | 2.31 | 0.00 | *G6PC3* | homozygous | NM_138387.4:c.758G>A, NP_612396.1:p.Arg253His |
| 1250 | m | neutropenia | 1 | 4.39 | 0.00 | *ELANE* | heterozygous | NM_001972.2:c.598-3C>G |
| 1251 | m | neutropenia | 1 | 6.97 | 0.00 | *ELANE* | heterozygous | NM_001972.2:c.598-3C>G |
| 1252 | m | neutropenia,neutropenia,after_BMT_cn | 3 | 1.66 | 0.52 | *ELANE* | heterozygous | NM_001972:c.640G>A, NP_001963.1:p.Gly214Arg |
| 1253 | f | neutropenia | 1 | 18.00 | 0.00 | *SBDS* | no_info | no_info |
| 1254 | f | neutropenia | 1 | 9.30 | 0.00 | *SRP54* | heterozygous | NP_115881.3:p.Thr117del |
| 1255 | f | neutropenia,neutropenia | 2 | 9.65 | 1.14 | *neg_exome* | | - |
| 1256 | m | neutropenia,neutropenia | 2 | 29.48 | 1.23 | *SBDS* | compound heterozygous | NM_016038.4:c.183_184delinsCT, NM_016038.4:258+2T>C |
| 1257 | f | neutropenia | 1 | 37.68 | 0.00 | *ELANE* | heterozygous | NP_001963.1:p.Cys223* |
| 1258 | f | neutropenia,neutropenia | 2 | 5.79 | 1.02 | *SBDS* | compound heterozygous | NM_016038.4:258+2T>C, deletion of exon 5 in SBDS |
| 1259 | m | neutropenia,neutropenia | 2 | 8.73 | 1.60 | *SBDS* | no_info | no_info |
| 1260 | f | neutropenia | 1 | 6.57 | 0.00 | *SBDS* | no_info | NM_016038.4:c.183_184delinsCT, NM_016038.4:258+2T>C |
| 1261 | f | neutropenia,neutropenia | 2 | 11.18 | 1.17 | *ELANE* | heterozygous | NM_001972.2:c.386_388delTCA |
| 1262 | m | neutropenia | 1 | 7.26 | 0.00 | *SBDS* | no_info | no_info |
| 1263 | f | neutropenia | 1 | 26.53 | 0.00 | *SBDS* | compound heterozygous | NM_016038.4:258+2T>C, NM_016038.4:c.460-1G>A |
| 1264 | m | neutropenia | 1 | 10.42 | 0.00 | *ELANE* | homozygous | NP_001963.1:p.Arg220Gln |
| 1265 | f | neutropenia | 1 | 17.91 | 0.00 | *JAGN1* | homozygous | NM_032492.3:c.63G>T, NP_115881.3:p.Glu21Asp |
| 1266 | f | neutropenia | 1 | 6.96 | 0.00 | *SBDS* | compound heterozygous | NM_016038.4:258+2T>C, deletion of exon 2 in SBDS |
| 1267 | m | neutropenia,neutropenia | 2 | 27.99 | 1.17 | *SBDS* | no_info | no_info |
| 1268 | m | neutropenia | 1 | 4.91 | 0.00 | *HAX1* | no_info | no_info |
| 1269 | f | neutropenia | 1 | 12.97 | 0.00 | *HAX1* | no_info | no_info |
| 1270 | f | neutropenia,neutropenia | 2 | 13.84 | 0.66 | *neg_exome* | | - |
| 1271 | m | neutropenia,neutropenia | 2 | 14.98 | 1.02 | *SRP54* | no_info | no_info |

| | | | | | | | | |
|---|---|---|---|---|---|---|---|---|
| 1272 | m | neutropenia | 1 | 1.87 | 0.00 | *ELANE* | heterozygous | NP_001963.1:p.Ser126Leu |
| 1273 | f | neutropenia | 1 | 1.96 | 0.00 | *ELANE* | heterozygous | NP_001963.1:p.Leu152Pro |
| 1274 | f | neutropenia | 1 | 8.67 | 0.00 | *CXCR4* | no_info | no_info |
| 1275 | f | neutropenia | 1 | 4.96 | 0.00 | *ELANE* | heterozygous | NP_001963.1:p.Gly214Glu |
| 1276 | m | neutropenia,neutropenia | 2 | 12.87 | 1.26 | *SRP54* | no_info | no_info |
| 1277 | m | neutropenia,neutropenia | 2 | 13.36 | 1.03 | *SBDS* | compound heterozygous | NM_016038.4:c.183_184delinsCT, NM_016038.4:258+2T>C |
| 1278 | m | neutropenia | 1 | 8.52 | 0.00 | *TAFAZZIN* | no_info | no_info |
| 1279 | f | neutropenia | 1 | 0.88 | 0.00 | *CLPB* | no_info | no_info |
| 1280 | m | neutropenia | 1 | 13.55 | 0.00 | *G6PC3* | no_info | no_info |
| 1281 | f | neutropenia | 1 | 18.23 | 0.00 | *G6PC3* | no_info | no_info |
| 1282 | m | neutropenia | 1 | 3.39 | 0.00 | *SBDS* | no_info | no_info |
| 1283 | f | neutropenia | 1 | 5.56 | 0.00 | *ELANE* | heterozygous | NM_001972.2:c.659G>A, NP_001963.1:Arg220Gln |
| 1284 | m | neutropenia | 1 | 0.94 | 0.00 | *ELANE* | heterozygous | NP_001963.1:p.Cys71Tyr |
| 1285 | m | neutropenia | 1 | 1.25 | 0.00 | *ELANE* | heterozygous | NM_001972.2:c.580+1G>A, NP_001963.1:p.Val190_Phe199del |
| 1286 | f | neutropenia | 1 | 8.89 | 0.00 | *SBDS* | no_info | no_info |
| 1287 | f | neutropenia | 1 | 9.64 | 0.00 | *ELANE* | heterozygous | NM_001972.2:c.301G>A, NP_001963.1:p.Val101Met |
| 1288 | f | neutropenia | 1 | 1.16 | 0.00 | *neg_exome* | | - |
| 1289 | f | neutropenia | 1 | 7.48 | 0.00 | *SBDS* | compound heterozygous | NM_016038.4:258+2T>C, NM_016038.4:c.18delC, NP_057122.2:p.Thr7Profs*6 |
| 1290 | m | neutropenia | 1 | 25.72 | 0.00 | *SBDS* | no_info | no_info |
| 1291 | m | neutropenia,neutropenia,neutropenia | 3 | 13.49 | 8.51 | *SBDS* | no_info | no_info |
| 1292 | m | neutropenia,neutropenia | 2 | 27.54 | 0.56 | *SRP54* | no_info | no_info |
| 1293 | m | neutropenia | 1 | 9.13 | 0.00 | *ELANE* | heterozygous | NM_001972.4:c.641G>T, NP_001963.1:p.Gly214Val |
| 1294 | m | neutropenia | 1 | 10.23 | 0.00 | *HAX1* | no_info | no_info |
| 1295 | m | neutropenia | 1 | 12.11 | 0.00 | *SBDS* | no_info | no_info |
| 1296 | m | neutropenia | 1 | 9.65 | 0.00 | *SBDS* | no_info | no_info |
| 1297 | f | neutropenia | 1 | 3.42 | 0.00 | *ELANE* | no_info | no_info |
| 1298 | f | neutropenia | 1 | 3.18 | 0.00 | *ELANE* | | NM_001972.2:c.377C>T, NP_001963.1:p.Ser126Leu |
| 1299 | m | neutropenia | 1 | 5.14 | 0.00 | *neg_exome* | | - |
| 1300 | m | neutropenia | 1 | 5.99 | 0.00 | *neg_exome* | | - |
| 1301 | m | neutropenia | 1 | 30.25 | 0.00 | *neg_exome* | no_info | no_info |
| 1302 | m | neutropenia | 1 | 14.47 | 0.00 | *ELANE* | heterozygous | NP_001963.1:p.Phe43Leu |
| 1303 | f | neutropenia | 1 | 7.71 | 0.00 | *HAX1* | homozygous | NP_006109.2:p.Trp44* |
| 1304 | m | neutropenia | 1 | 9.22 | 0.00 | *HAX1* | homozygous | NP_006109.2:p.Trp44* |
| 1305 | f | neutropenia | 1 | 32.95 | 0.00 | *ELANE* | no_info | no_info |
| 1306 | f | neutropenia | 1 | 21.36 | 0.00 | *SBDS* | no_info | no_info |
| 1307 | f | neutropenia | 1 | 5.55 | 0.00 | *ELANE* | no_info | no_info |
| 1308 | m | neutropenia | 1 | 6.11 | 0.00 | *HAX1* | homozygous | NP_006109.2:p.Trp44* |

| | | | | | | | | |
|---|---|---|---|---|---|---|---|---|
| 1309 | f | neutropenia | 1 | 19.12 | 0.00 | *SBDS* | no_info | no_info |
| 1310 | f | neutropenia | 1 | 35.56 | 0.00 | *SBDS* | no_info | no_info |
| 1311 | f | neutropenia | 1 | 24.52 | 0.00 | *SBDS* | compound heterozygous | NM_016038.4:c.183_184delinsCT, NM_016038.4:258+2T>C |
| 1312 | f | neutropenia | 1 | 28.94 | 0.00 | *SBDS* | compound heterozygous | NM_016038.4:c.183_184delinsCT, NM_016038.4:258+2T>C |
| 1313 | m | neutropenia | 1 | 23.57 | 0.00 | *SBDS* | compound heterozygous | NM_016038.4:c.183_184delinsCT, NM_016038.4:258+2T>C |
| 1314 | m | leukemia | 1 | 0.91 | 0.00 | *ELANE* | heterozygous | NP_001963.1:p.Asp230Metfs*10 |
| 1315 | m | progressive,MDS | 2 | 5.37 | 4.56 | *SBDS* | compound heterozygous | NM_016038.4:258+2T>C, NM_016038.4:c.164C>G, NP_057122.2:p.Ser55* |
| 1316 | f | neutropenia | 1 | 0.90 | 0.00 | *SRP54* | heterozygous | NM_003136.4:c.349_351del, NP_003127.1:p.Thr117del |
| 1317 | m | neutropenia | 1 | 2.96 | 0.00 | *SBDS* | compound | NM_016038.4:c.183_184delinsCT, NM_016038.4:258+2T>C |
| 1318 | f | neutropenia | 1 | 0.91 | 0.00 | *ELANE* | no_info | no_info |
| 1319 | f | neutropenia | 1 | 5.44 | 0.00 | *ELANE* | heterozygous | NM_001972.2:c.580C>T, NP_001963.1:p.Gln194* |

**Supplementary table 3. List of genes/gene regions covered by capture panel version 1.**

| | Gene names |
|---|---|
| Sequencing of entire coding gene sequence (n=31) | *ASXL1, BCOR, BCORL1, CBL, CEBPA, CSF3R, DNMT3A, EP300, EZH2, FLT3, IDH1, IDH2, JAK2, KDM6A, KRAS, NPM1, NRAS, PHF6, PTPN11, RAD21, RUNX1, SETBP1, SF1, SF3B1, SRSF2, STAG2, SUZ12, TET2, TP53, U2AF1, ZRSR2* |
| Sequencing of "hot spot" regions (n=340). The regions (mutated in at least 2-3 cancer patients) were retrieved from whole exome/genome sequencing datasets of myeloid neoplasm samples (n=2613) available on https://www.cbioportal.org. Genomic coordinates are available on request. | *ABL1, ACAN, ACSL3, AFF1, AFF4, ALK, AMER1, ANKRD11, APC, ARHGAP35, ARHGEF28, ARID1A, ARID1B, ARID2, ARID4A, ARID4B, ASXL2, ATM, ATP1A1, ATP2B3, ATRX, AURKB, AXL, BABAM1, BCL9, BPIFC, BRCA2, BRCC3, BRD4, BRINP3, CACNA1B, CACNA1D, CACNA1E, CACNA2D3, CAD, CALR, CAMTA1, CBFB, CBLB, CCND2, CCND3, CD70, CD74, CDC42, CDH11, CENPA, CHD2, CHD4, CHEK2, CHIC2, CHN1, CIC, CIITA, CLTCL1, CMYA5, CNOT3, COL1A1, COL2A1, CPS1, CREB3L1, CREBBP, CROCC, CSDE1, CSMD1, CSMD2, CSMD3, CTCF, CUL3, CUX1, CYLD, DAXX, DCLK1, DCTN1, DDR2, DDX41, DICER1, DIS3, DNAH3, DNAH9, DNM2, DNMT1, DNMT3B, DOT1L, DROSHA, DSCAM, DYSF, EED, EGFR, EGR1, ELF4, ELL, ELN, EML4, EP400, EPHA3, EPHA5, EPHA7, EPHB1, ERBB2, ERBB3, ERBB4, ERG, ESCO2, ETNK1, ETV1, ETV6, EWSR1, EZH1, EZR, FAF1, FANCA, FANCC, FAT1, FAT4, FBXO11, FCGBP, FLT1, FOXP1, FREM2, FRYL, GATA1, GATA2, GATA3, GIGYF2, GLI1, GNAS, GNB1, GPC3, GRIK2, GRIK4, GRIN2A, GRIN2B, GRM3, HDAC4, HECW1, HERPUD1, HIST1H1C, HIST1H1E, HIST1H3C, HIST1H3D, HLA-B, HLA-C, HNRNPK, HOOK3, HOXB13, HSD3B1, IKBKB, IKZF1, IL6ST, INPP4A, INPP4B, INPP5D, INPPL1, IRS2, IRS4, JAK3, JARID2, KAT6A, KAT6B, KBTBD4, KDM2B, KDM5C, KDR, KIF5B, KIT, KMT2A, KMT2B, KMT2C, KMT2D, KSR2, LRIG3, LRP1B, LRP5, LRP6, LRRK2, MAP2, MAP3K13, MAP3K7, MAPK1, MAX, MDC1, MDM2, MDM4, MED12, MEF2B, MGA, MKI67, MN1, MPL, MSH2, MSH3, MST1, MST1R, MTA2, MTOR, MXRA5, MYB, MYC, MYCBP2, MYD88, MYH11, MYO18A, MYO5A, NADK, NCOA2, NCOA3, NCOR1, NF1, NFATC2, NFE2, NIN, NONO, NOTCH1, NOTCH2, NOTCH3, NOTCH4, NR4A3, NSD1, NTRK3, NUMA1, NUP214, NUP93, NUP98, PAPPA2, PARP2, PASK, PAX3, PAX5, PBRM1, PBX1, PC, PCLO, PDCD11, PDE4DIP, PDGFRA, PDGFRB, PDS5B, PGBD5, PHIP, PIK3C2B, PIK3C3, PIK3CA, PIK3CB, PIK3R1, PKD1L2, PLCE1, PLCG1, PMS2, POLE, POLQ, POU5F1, PPFIBP1, PPM1D, PPP2R2A, PPP6C, PRDM1, PRDM16, PREX2, PRKD1, PSIP1, PTCH1, PTEN, PTPRC, PTPRD, PTPRS, PTPRT, RABEP1, RAC2, RAD51, RANBP2, RAP1GDS1, RASA1, RELN, RIT1, RNF213, ROBO1, ROS1, RPTOR, RRAS2, RUNX1T1, SALL4, SEC31A, SERPINB4, SETD1B, SETD2, SETD5, SH2B3, SHQ1, SI, SMARCA2, SMC1A, SMC3, SMG1, SNCAIP, SNX29, SOCS1, SOS1, SPEN, STAG1, STAT2, STAT4, STAT5B, STRN, SYK, TAF1, TAP2, TBL1XR1, TCF12, TCF7L2, TCL1A, TEK, TENM2, TET1, TGFBR2, TLL2, TLX3, TNC, TNFRSF11A, TOP1, TPR, TRAF7, TRIM24, TRIM33, TRIP11, TRPM3, TRRAP, TSHR, TUSC3, TYRO3, U2AF2, UBR5, UPF1, USP6, WIF1, WRN, WT1, ZBTB33, ZBTB7A, ZEB2, ZFHX3, ZFP36L2, ZNF687* |

**Supplementary table 4. List of genes/gene regions covered by capture panel version 2.**

| | Gene names |
|---|---|
| Sequencing of the entire coding gene sequence (n = 29) | *ASXL1, ASXL2, BCOR, BCORL1, CBL, CEBPA, CSF3R, DDX41, DNMT3A, ETV6, EZH2, KIT, KMT2D, KRAS, NF1, NPM1, NRAS, PHF6, PTPN11, RAD21, RUNX1, SETBP1, SRSF2, STAG2, TET2, TP53, U2AF1, WT1, ZRSR2* |
| Sequencing of hotspot regions in leukemia-associated genes (n = 16). The DNA positions (mutated in at least 2-3 cancer patients) were selected using the whole exome and whole genome sequencing records of 9863 leukemia and MDS samples (9569 patients) available on https://www.cbioportal.org. A detailed description of datasets is available at https://bit.ly/3XX8iKv. Genomic coordinates are available on request. | *ATM, CTCF, EP300, ETNK1, FLT3, GATA1, GATA2, IDH1, IDH2, JAK2, MPL, SF3B1, SMC1A, SMC3, SRCAP, SUZ12* |

**Supplementary table 5. Filtering criteria for somatic mutation detection in sequencing data.**

| Filter name and description | CN, SDS | CN/SDS prior MDS/AML | SDS-, CN-MDS/AML |
|---|---|---|---|
| Mutect2 standard somatic variants filter | + | + | + |
| Mutect2 gnomAD based germline variants filter | + | + | + |
| Filtering out variants detected by Mutect2 in Panel of normals (PON) | + | + | + |
| Keep non-synonymous alterations[1,2] | + | + | + |
| Remove variants with population allele frequency >1e-04[3] | + | + | + |
| Remove variants with allele frequency > 0.26[3,4,5] | + | + | - |

[1]Funcotator variant annotations: Frame_Shift_Del, Frame_Shift_Ins, Splice_Site, Translation_Start_Site, Nonsense_Mutation, Nonstop_Mutation, In_Frame_Del, In_Frame_Ins, Missense_Mutation; [2]Non-coding and synonymous variants were included in the analysis of longitudinal samples from CN-AML patients (**Figure S1D, S1E, S5A**); [3]Except variants from ClinVar interpretation: Likely pathogenic, Pathogenic, Pathogenic/Likely pathogenic, risk factor, drug response; [4]Except manually curated samples/variants; [5]Except nonsense mutations and frameshift/inframe indels.

### Supplementary table 6. Recurrent cytogenetic aberrations in SDS- and CN-MDS/AML.

| Patient ID | Genetic diagnosis | Age at MDS diagnosis, years | MDS type | Leukemia onset age, years | Leukemia phenotype | % of blasts in PB at leukemia, (MDS) | % of blasts in BM at leukemia, (MDS) | Loss of chr 7 | Trisomy chr 21 | Additional Cytogenetic Findings |
|---|---|---|---|---|---|---|---|---|---|---|
| 1007 | *ELANE-CN* | | | 21.17 | AML M2 | 29.5 | 49 | no | add(21)(q22) | Hyperdiploidy (49-50 chromosomes), +8, +12, +20 |
| 1213 | *ELANE-CN* | | | 55.03 | pro-B-ALL | 16 | 60 | no | yes | del(5)(q14q34), rearrangements at 3q26, 8q22, and 11q23 |
| 1010 | *ELANE-CN* | | | 16.78 | AML M4 | 35 | 15 | yes | no | Tetrasomy 8, tetrasomy 18 |
| 1015 | *ELANE-CN* | 2.51 | | | | | | del q7 | no | |
| 1224 | *ELANE-CN* | | | 5.34 | AML | | | N/A | N/A | N/A |
| 1006 | *ELANE-CyN* | | | 53.06 | atypical CML | | | no | no | Normal karyotype (46,XY) |
| 1178 | *ELANE-CyN* | | | 27.85 | AML | | 20-25 | del(7)(q21q32) | tidic(21)(p11) | Complex: t(5;7), -16, add(2q) |
| 1013 | *JAGN1* | | | 19.14 | AML M2 | 19.5 | 25 | yes | no | |
| 1054 | *HAX1* | 0.98 | | | | 0 | 5 | no | no | Normal karyotype (46,XY) |
| 1014 | *ELANE-CN* | | | 2.93 | AML/B-ALL | 47 | 64 | no | Trisomy 21; 21q22(AML1x 3-4) | - |
| 1009 | *HAX1* | 15.91 | RAEB | 15.98 | CMML | 1 | 20 | no | no | Normal karyotype (46,XY) |
| 1016 | *SDS* | 14.35 | RAEB | | | 0 | 4 | yes | no | t(2;5)(q32;q11), presence of a ring chromosome (+r) |
| 1008 | *HAX1* | 11.32 | RAEB-T | 11.32 | | 13 | 22.5 | no | no | Normal karyotype (46,XY) |
| 1011 | *HAX1* | 4.51 | RAEB-T | 4.56 | AML | 5.5 | 22, (11.5) | yes | no | - |
| 1148 | *WAS* | | | 16.42 | AML | 0 | 30 | yes | no | Complex clonal evolution: del(18)(q22), del(13)(q13q33), and a derivative translocation der(6)t(3;6)(q13;p24). |
| 1004 | *HAX1* | 6.85 | RAEB-T | 6.85 | | 18 | 32.5 | no | no | Normal karyotype (46,XY) |
| 1191 | *HAX1* | 9.93 | | 10.06 | AML | 0 | 49, (1) | no | no | del(9)(q21q33) |
| 1093 | *SBDS* | | | 24.44 | | | | N/A | N/A | N/A |
| 1199 | *SLC37A4* | | | 10.74 | AML M2 | 25 | 20 | no | yes | - |
| 1001 | *ELANE-CyN* | | | 38.36 | common B-ALL | 2 | 70 | yes | Tetrasomy 21, trisomy 21 | Trisomy 22, del(5)(q13q33), +mar |
| 1002 | *ELANE-CyN* | 17.68 | | 17.95 | AML M2 | 22, (0) | | yes | yes | - |
| 1211 | *SBDS* | 11.09 | | | | | 33, (4) | yes | N/A | N/A |
| 1166 | *SBDS* | N/A | N/A | N/A | N/A | | | N/A | N/A | N/A |

| | | | | | | | | | | |
|---|---|---|---|---|---|---|---|---|---|---|
| 1314 | *ELANE* | N/A | N/A | 0.91 | N/A | | | yes | N/A | N/A |
| 1226 | *HAX1* | | | 14.85 | AML | 42 | 20 | yes | no | t(2;3)(q22;q27) involving a break in the EVI1 gene at 3q26. |
| 1200 | *ELANE-CN* | | | 10.60 | AML M4 | 80 | 79 | no | no | Trisomy 12, add(7)(q21) |
| 1005 | *ELANE-CN* | 22.38 | RAEB-T | 23.20 | AML M1 | 12-23 | | yes | yes | - |
| 1003 | *ELANE* | | | 18.51 | AML M4 | | | no | no | t(9;11) |
| 1162 | *WAS* | 20.30 | RAEB | 20.78 | AML | | | no | no | del(2)(q35), add(7)(q21) |
| 1160 | *ELANE* | 6.95 | | 7.60 | AML | | 11 | no | no | Normal karyotype (46,XY) |
| 1158 | *ELANE* | | | 14.96 | AML M5 | | 30 | yes | no | - |
| 1150 | *ELANE* | | | 55.42 | AML | | 70 | no | yes | der(4)t(1:4)(q21:q25), del(5)(q13q31), add(7)(q22) |
| 1159 | *ELANE* | | | 17.70 | AML M5 | | 25 | del(7)(q22q36) | yes | add(7)(q22), der(1)t(1,3), del(5)(q31q35) |
| 1202 | *SBDS* | N/A | N/A | N/A | N/A | | | N/A | N/A | N/A |
| 1012 | *WAS* | | | 7.13 | AML M0 | | | N/A | N/A | N/A |
| 1161 | *SBDS* | 12.49 | | | | | | N/A | N/A | N/A |
| 1315 | *SBDS* | | | 5.11 | AML | | | N/A | N/A | N/A |
